# Pursuing the Mechanisms Underlying Alcohol-Induced Changes in the Ghrelin System: New Insights from Preclinical and Clinical Investigations

**DOI:** 10.1101/2020.07.30.228494

**Authors:** Mehdi Farokhnia, Sara L. Deschaine, Adriana Gregory-Flores, Lia J. Zallar, Zhi-Bing You, Hui Sun, Deon M. Harvey, Renata C.N. Marchette, Brendan J. Tunstall, Bharath K. Mani, Jacob E. Moose, Mary R. Lee, Eliot Gardner, Fatemeh Akhlaghi, Marisa Roberto, James L. Hougland, Jeffrey M. Zigman, George F. Koob, Leandro F. Vendruscolo, Lorenzo Leggio

## Abstract

Ghrelin is a gastric-derived peptide hormone with demonstrated impact on alcohol intake and craving, but the reverse side of this bidirectional link, i.e., the effects of alcohol on the ghrelin system, remains to be fully established. To characterize the downstream effects of alcohol on the ghrelin system, we examined the following: (1) plasma ghrelin levels across four human laboratory alcohol administration experiments with non-treatment seeking, heavy-drinking participants, (2) expression of ghrelin, ghrelin receptor, and ghrelin-O-acyltransferase (GOAT) genes *(GHRL, GHSR,* and *MBOAT4*, respectively) in human *post-mortem* brain tissue from individuals with alcohol use disorder (AUD) *vs.* controls, (3) plasma ghrelin levels in *Ghsr* knockout and wild-type rats following intraperitoneal (i.p.) ethanol administration, (4) effect of ethanol on ghrelin secretion from gastric mucosa cells *ex vivo* and GOAT enzymatic activity *in vitro,* and (5) plasma ghrelin levels in rats following i.p. ethanol administration *vs*. an iso-caloric sucrose solution. Peripheral acyl- and total ghrelin levels significantly decreased following acute ethanol administration in humans. No difference in *GHRL, GHSR,* and *MBOAT4* mRNA expression in the brain was observed between AUD *vs.* control *post-mortem* samples. In rats, acyl-ghrelin levels significantly decreased following i.p. ethanol administration in both genotype groups *(Ghsr* knockout and wild-type), while des-acyl-ghrelin was not affected by ethanol. No effect of ethanol was observed *ex vivo* on ghrelin secretion from gastric mucosa cells or *in vitro* on GOAT acylation activity. Lastly, we observed different effects of i.p. ethanol and sucrose solution on acyl- and des-acyl-ghrelin in rats despite administering amounts with equivalent caloric value. Ethanol acutely decreases peripheral ghrelin concentrations in humans and rats, and our findings suggest that this effect does not occur through interaction with ghrelin-secreting gastric mucosal cells, the ghrelin receptor, or the GOAT enzyme. Moreover, this effect does not appear to be proportional to caloric load. Our findings, therefore, suggest that ethanol does not suppress circulating ghrelin through direct interaction with the ghrelin system, or in proportion to the caloric value of alcohol, and may differentially affect ghrelin acylation and ghrelin peptide secretion.

## 1. Introductio

Alcohol use disorder (AUD) is a chronic relapsing disease characterized by excessive consumption of alcohol to an extent that causes significant harm to the affected individual’s health and overall quality of life. According to the 2018 National Survey on Drug Use and Health, 5.8% of individuals aged 18 and older in the United States had AUD in the past year, and an estimated 88,000 annual deaths are alcohol-related [1, 2]. Still, only three Food and Drug Administration (FDA)-approved medications are available for treatment of AUD, highlighting a significant need to develop novel pharmacotherapies for AUD. One such therapeutic strategy is based on the notion that harmful alcohol consumption can be alleviated by pharmacologically manipulating endocrine pathways that control both homeostatic and hedonic feeding, as well as stress-related pathways and reward processing [3–5]. Indeed, the orexigenic peptide ghrelin is one hormone that has been shown to play a role in alcohol-related behavior across numerous studies [6–8].

Ghrelin is a 28 amino acid hormone secreted primarily from P/D1 cells (X/A-like cells in rodents) located in the oxyntic glands of the fundus portion of the stomach. Encoded by the ghrelin gene *(GHRL),* ghrelin is post-translationally formed by cleavage of the 117 amino acid preproghrelin into proghrelin, which can then be acylated at the serine-3 residue by the membrane-bound enzyme, ghrelin O-acyltransferase (GOAT) [9–12]. Acylated proghrelin is then cleaved to form acyl-ghrelin – the endogenous ligand of the growth hormone secretagogue receptor 1a (GHSR1a). Acylation of ghrelin is essential for binding to GHSR1a, both centrally and peripherally, and mediates orexigenic effects [9, 13, 14]. Much research over the past decade demonstrates that the ghrelin system has a complex biology, due to several factors: (1) circulating acyl-ghrelin in can be de-acylated by plasma esterases to des-acyl-ghrelin [15]; (2) GOAT can acylate ghrelin in target tissues, both in the central nervous system and the periphery [16–18]; (3) plasma anti-ghrelin immunoglobulin Gs (IgGs) may bind and protect ghrelin from degradation in circulation [19]; (4) des-acyl-ghrelin may have effects seemingly opposite to acyl-ghrelin through GHSR1a-independent mechanisms [20]; and (5) GHSR1a has high constitutive, ligand-independent activity [21, 22]. Moreover, an endogenous antagonist/inverse agonist for GHSR1a, known as liver-expressed antimicrobial peptide-2 (LEAP-2), was recently identified [23–25]. These different components of the ghrelin system help regulate and balance acyl-ghrelin’s important effects on energy homeostasis to ensure survival of the organism [26].

Central signaling of the gastric-derived acyl-ghrelin occurs through activation of GHSR1a expressed on vagal afferent neurons in the stomach, as well as by acyl-ghrelin crossing the blood-brain barrier and binding to GHSR1a in the brain [27]. Acyl-ghrelin’s central orexigenic signaling occurs directly through GHSR1a expressed on hypothalamic neuropeptide Y and agouti-related, peptide-expressing neurons in the arcuate nucleus, and indirectly through activation of the lateral hypothalamus, hippocampus, amygdala, ventral tegmental area (VTA), and other regions [27]. These brain regions communicate with origins and terminal regions of the mesolimbic dopamine system, which affects motivational components of behaviors, including hedonic feeding and drug and alcohol seeking. Indeed, administration of acyl-ghrelin into the brain’s reward circuitry increases extracellular dopamine via GHSR1a, located in the mesolimbic pathway [28–35], and stimulates food intake [36, 37]. By communicating with these regions, the ghrelin system can regulate homeostatic and hedonic drives governing food-seeking behavior that seem to similarly affect alcohol-seeking behavior. In rodents, central or systemic ghrelin administration increases alcohol intake, whereas antagonism of GHSR1a and knockout of *Ghrl* or *Ghsr* decreases alcohol preference and consumption and blunts both conditioned place preference and dopamine release in the nucleus accumbens (NAc) induced by alcohol [38–43]. Moreover, higher peripheral ghrelin concentrations are positively correlated with alcohol craving and risk of relapse in humans [40, 44–48], and exogenous ghrelin administration increases cue-induced craving [45] and intravenous self-administration of alcohol in heavy-drinking individuals with alcohol dependence [49]. Collectively, these studies demonstrate a clear relationship between ghrelin and alcohol-related behavior, wherein ghrelin appears to potentiate alcohol seeking and consumption, and partly regulate its reinforcing effects.

To obtain a better understanding of the crosstalk between ghrelin and alcohol, further research into how alcohol affects the ghrelin system is critically needed. To date, only a few studies have examined the effect of alcohol on peripheral ghrelin concentration. In rodents, alcohol acutely decreased both acyl-ghrelin and total-ghrelin concentration in plasma [50, 51], and in humans, acute administration of alcohol decreased plasma ghrelin concentrations [52–57]. Amongst individuals with alcohol dependence, abstainers had higher peripheral ghrelin concentrations compared to current drinkers [44, 47, 48, 58–63]. Ghrelin concentrations were significantly lower in individuals with alcohol dependence compared to matched, non-dependent controls [44]. However, mean daily alcohol consumption over the past 12 months was positively correlated with plasma ghrelin concentrations among individuals without an AUD diagnosis [64]. Collectively, these studies suggest that acute and chronic exposure to alcohol differentially affect the ghrelin system, with effects from chronic alcohol use likely reflecting compensatory mechanisms dependent on the extent and duration of an individual’s alcohol use. Although the literature to date demonstrates an interplay between alcohol and ghrelin, it is unclear how these effects are occurring – whether through direct action on ghrelin-producing cells, modification of GOAT activity, and/or through other mechanism(s). The objective of the present body of work was to further probe the effect of alcohol on the ghrelin system and its underlying mechanisms through experiments performed in humans and rodents, *ex vivo* experiments, and *in vitro* experiments.

## 2. Results

### 2.1 Plasma Ghrelin Levels are Reduced after Alcohol Administration in Humans

We examined plasma ghrelin levels following intravenous or oral alcohol administration during four different human laboratory experiments in non-treatment-seeking, heavy-drinking individuals (**Table S1, Figure S1-4**). Each experiment was originally conducted as part of a parent study designed to evaluate the effects of a pharmacological intervention on alcohol-related outcomes [49, 65, 66]. Here, we performed secondary analyses for each experiment using data from the placebo groups of these parent studies. Each experiment had a unique design, with alcohol being administered orally or intravenously (IV) in both self-administration and fixed-dose schedules (four experiments in total) (see **Figure 1** and **Table S2** for details). We evaluated changes in acyl-ghrelin and total-ghrelin during each alcohol administration experiment, which allowed us to compare different routes of alcohol administration.

**Figure 1:**
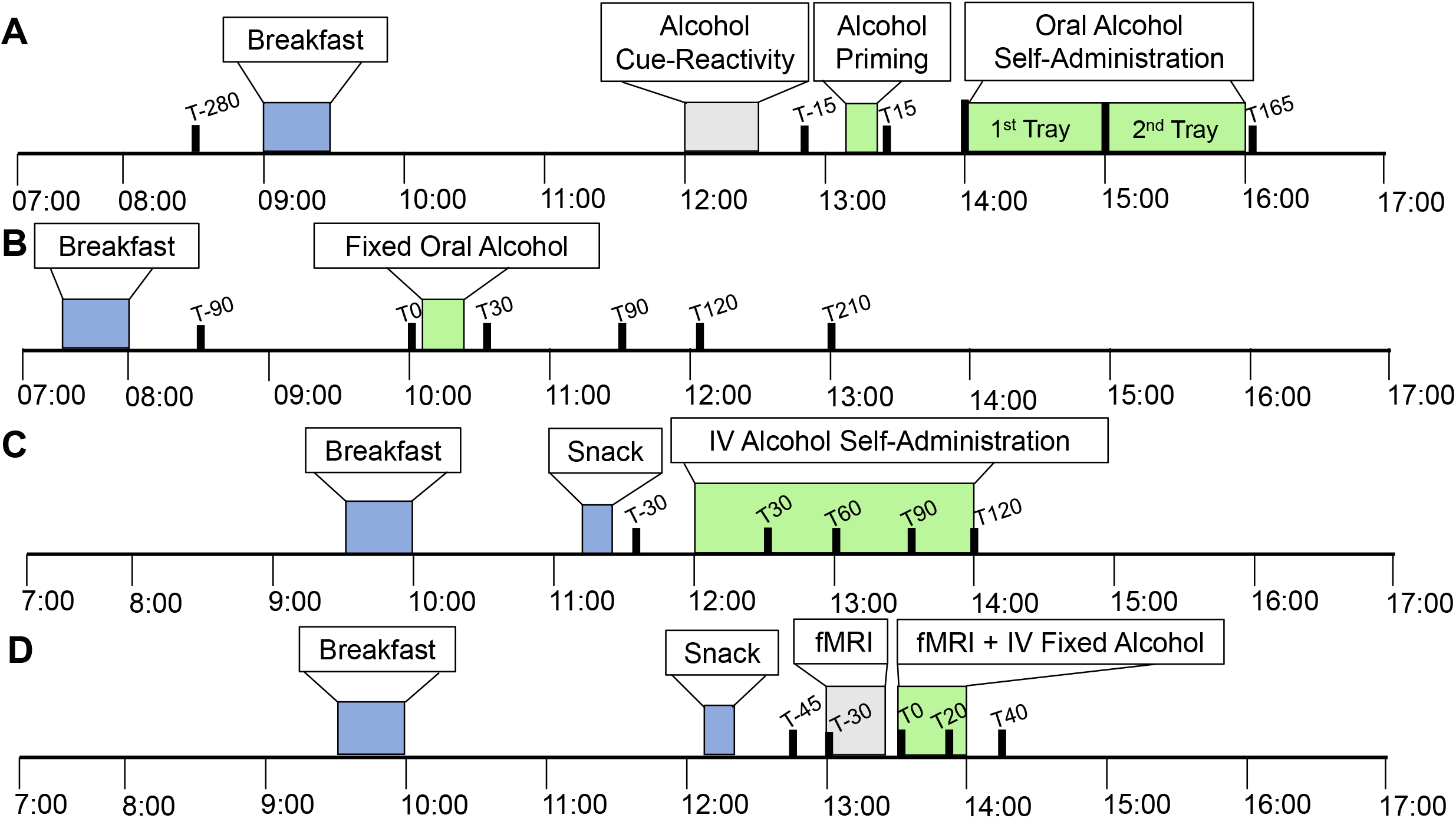
Schematic Overview of Human Laboratory Alcohol Administration Experiments. Overview of human laboratory experiments depicting meals (blue), duration of alcohol intake (green), and blood draw times (black). Other study procedures not involving alcohol administration are also outlined (gray) **1A:** Oral priming and alcohol self-administration experiment. Blood draw timepoints relative to 0 = alcohol administration at 13:15 are T-280, T-15, T15, T165 min **1B:** Fixed oral alcohol administration experiment. Blood draw timepoints relative to 0 = alcohol administration at 10:15 are T-90, T0, T30, T90, T120, and T210 min **1C:** Intravenous variable dose alcohol self-administration experiment. Blood draw timepoints relative to 0 = alcohol administration at 12:00 are T-30, T30, T60, T90, and T120 min **1D:** Fixed intravenous alcohol administration experiment. Blood draw timepoints relative to 0 = alcohol administration at 13:00 are T-45, T0, T20, T30, and T40 min.

Overall, alcohol administration led to a reduction in ghrelin levels, regardless of the route of ethanol administration, within a time period ranging from 45 – 165 min. Using linear mixed effects modeling, we found that acyl-ghrelin [F(3, 27.5) = 6.6, p = 0.002] and total ghrelin [F(3, 37.7) = 4.5, p = 0.009] were significantly reduced during the variable dose oral alcohol administration session (**Figure 2A**). Moreover, there was a significant reduction in acyl-ghrelin [F(5, 53.8) = 10.5, p < 0.001] and total ghrelin [F(5, 52) = 13.6, p < 0.001] during the fixed oral alcohol administration session (**Figure 2B**). Analysis of peripheral ghrelin during the variable dose IV alcohol administration session also revealed a significant reduction in both acyl-ghrelin [F(4,37.7) = 7.5, p < 0.001], and total ghrelin [Covariate: Gender, F(4, 38.7) = 5.6, p = 0.001] (**Figure 2C**). Lastly, we observed a reduction in acyl-ghrelin [F (4,19) = 2.0, p = 0.134) and total-ghrelin [F(4,17.1) = 2.7, p = 0.067] during the fixed IV alcohol administration session, but this change did not reach statistical significance (**Figure 2D**). Pairwise comparisons corrected for multiple testing were conducted between all timepoints for experiments where significant overall effects were found and are presented in **Figure 2**.

**Figure 2:**
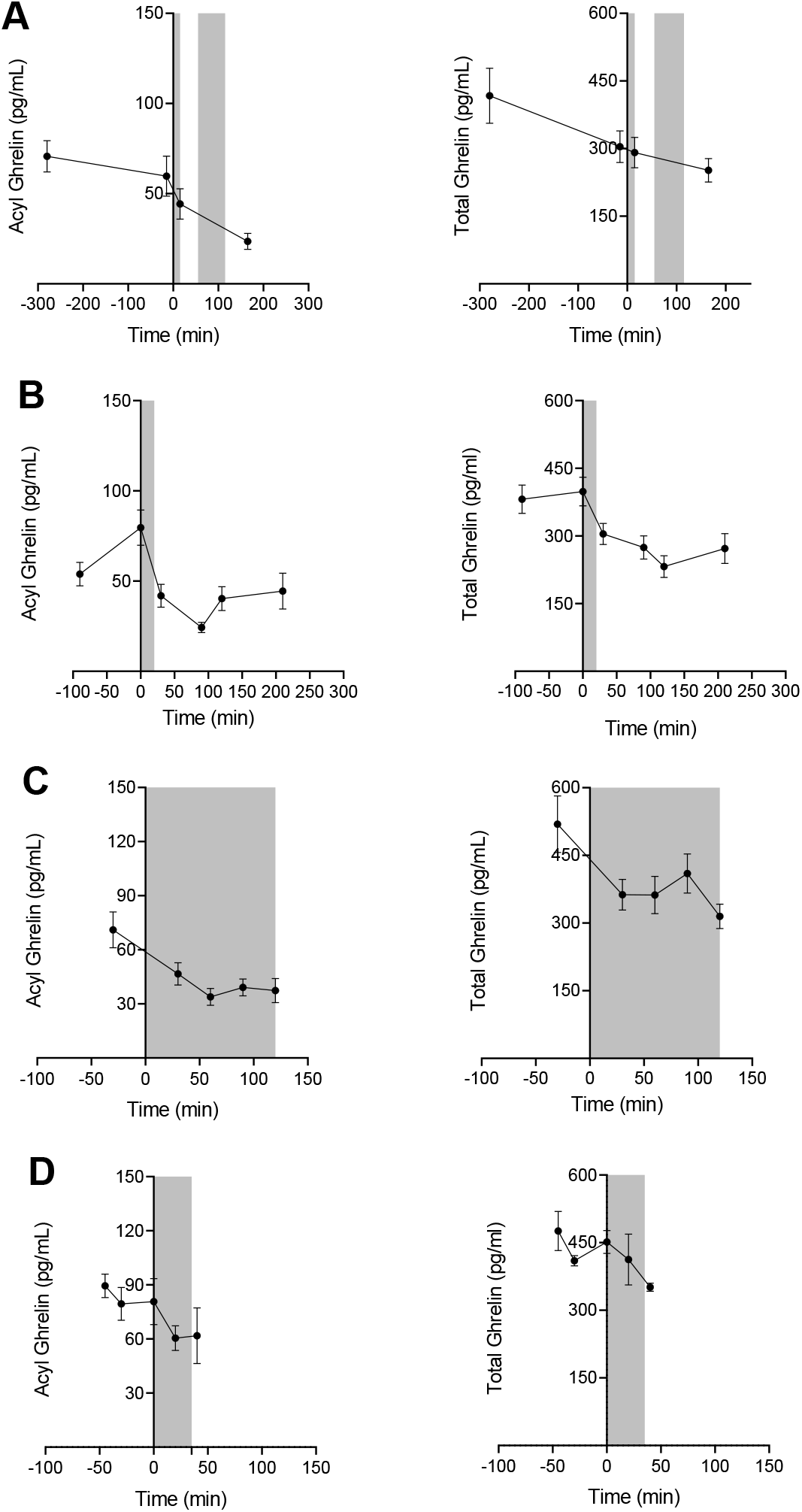
Effect of Alcohol Administration on Peripheral Ghrelin Levels in Humans. Plasma ghrelin levels over the course of different alcohol administration experiments in participants with heavy drinking. Gray zones indicate time periods of alcohol administration. For all data, 0 min = beginning of alcohol administration session. Data are presented as Mean ± SEM. **2A:** Oral variable-dose alcohol administration (Oral Priming and self-administration) analysis (N = 16); fixed effect of time (−280, −15, 15, 165) on acyl-ghrelin (Left: p < 0.002) and total ghrelin (Right: p = 0.009); pairwise comparisons: acyl-ghrelin (−280 *vs.* 165, −15 *vs.* 165; p < 0.05), total ghrelin (−280 *vs.* 165; p < 0.05). **2B:** Oral fixed-dose alcohol administration analysis (N = 12); fixed effect of time (−90, 0, 30, 90, 120, and 210 min) on acyl-ghrelin (Left: p < 0.0001) and total ghrelin (Right: p < 0.0001); pairwise comparisons: acyl-ghrelin (0 *vs.* −90; p < 0.05, 0 *vs.* 30, 90, 120, 210; p < 0.001), total ghrelin (0 *vs.* −90, 30, 210; p < 0.05, 0 *vs.* 120, p < 0.001) **2C:** IV variable-dose alcohol administration analysis (N = 11); fixed effect of time (−30, 30, 60, 90, and 120 min) on acyl-ghrelin (Left: p < 0.0001), and total ghrelin (Right: p < 0.0001); pairwise comparisons: acyl-ghrelin (−30 *vs.* 30; p < 0.05, −30 *vs.* 60, 90, 120; p < 0.001), total ghrelin (−30 *vs.* 30, 60; p < 0.05, −30 *vs.* 120; p < 0.01). **2D**: IV fixed-dose alcohol administration analysis (N = 6) evaluated a fixed effect of timepoint (−45, 0, 20, 30, and 40 min) on acyl-ghrelin (Left: p = *NS),* and total ghrelin (Right: p = *NS*). P values presented for pairwise comparisons are Bonferroni corrected.

### 2.2 *GHSR, GHRL,* and *MBOAT4* Expression Levels in Select Brain Regions Are Not Significantly Altered by Chronic Ethanol Consumption

We further examined the effect of alcohol on the ghrelin system by determining whether severe alcohol use disorder (AUD), as indexed by DSM-5 AUD, affected central expression levels of *GHRL, GHSR,* or the GOAT enzyme gene, *MBOAT4.* Fold changes of *GHSR* mRNA, *GHRL* mRNA, and *MBOAT4* mRNA expression levels in postmortem brain tissue [regions that have been shown to be affected by long term alcohol use (hippocampus, VTA, amygdala, prefrontal cortex (PFC) – superior frontal Brodmann area 12 and 13, and NAc] were compared between AUD individuals (N = 11) and controls (N = 15-16) (**Table 1**). Baseline characteristics of the sample are provided in **Table S3**. Using linear mixed effects modeling, we found no significant effect of group (AUD *vs.* control) on *GHRL* or *GHSR* expression in any of these brain regions tested (**Figure 3, Table 1**).

**Figure 3:**
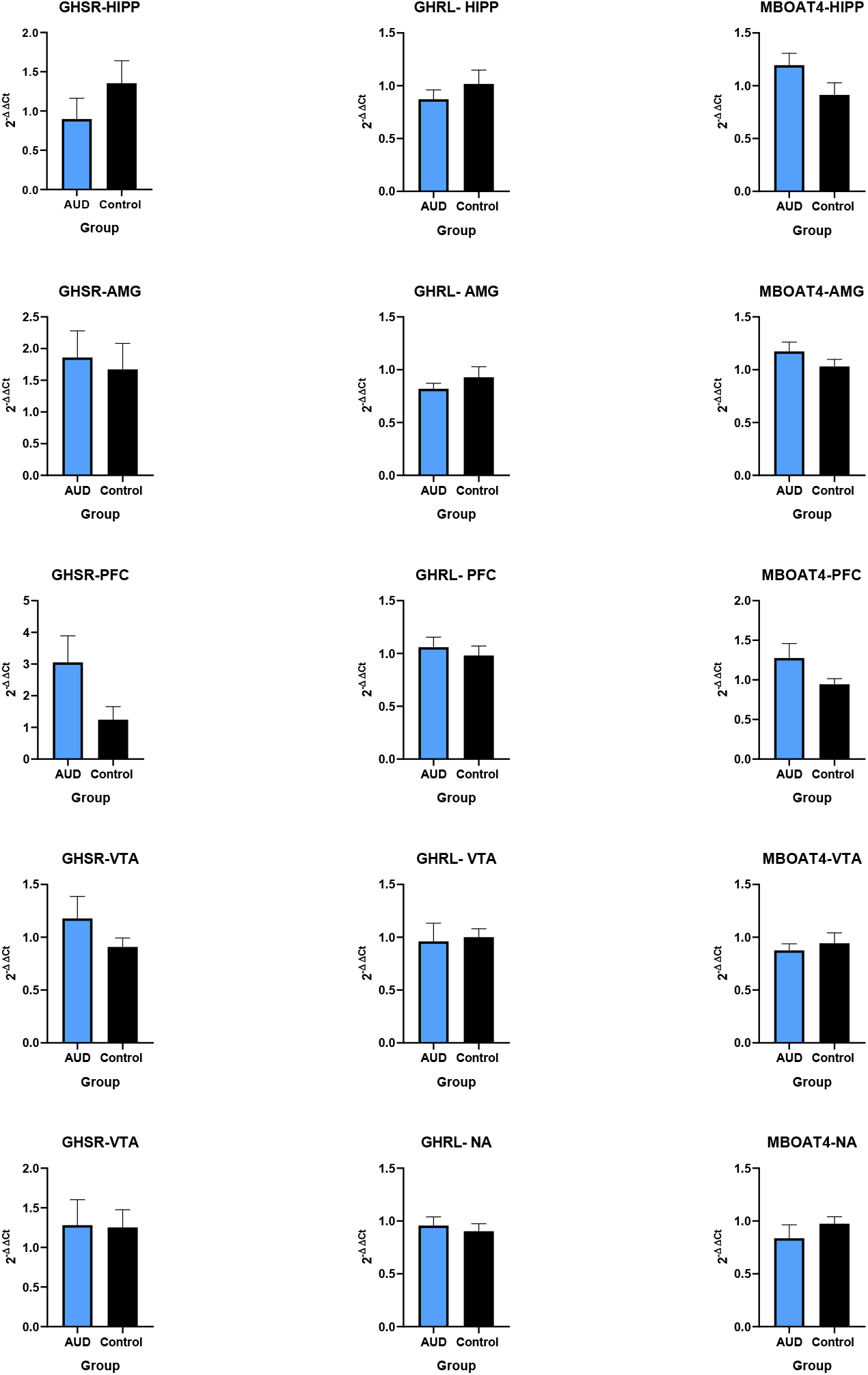
Central *Post Mortem* Expression of *GHSR, GHRL, and MBOAT4* in AUD Individuals and Controls. Fold expression of *GHSR, GHRL,* and *MBOAT4* in 5 selected brain regions examined in *post-mortem* brain tissue from individuals with AUD and controls. Fold expression change is expressed as 2^-ΔΔCt^ where ΔΔCt is the difference in ΔCt between AUD and control samples and ΔCt is difference in cycle threshold (Ct) for the gene of interest – *GADPH* in the same sample. No regions are significantly different from each other after controlling for multiple comparisons. AMG = amygdala, AUD = alcohol use disorder, GHRL = growth hormone receptor ligand (ghrelin gene), GHSR = growth hormone secretagogue receptor (ghrelin receptor gene), HIPP = hippocampus, MBOAT4 = membrane bound o-acyl transferase 4 (GOAT gene), NA = nucleus accumbens, PFC = prefrontal cortex, VTA = ventral tegmental area.

**Table 1:**
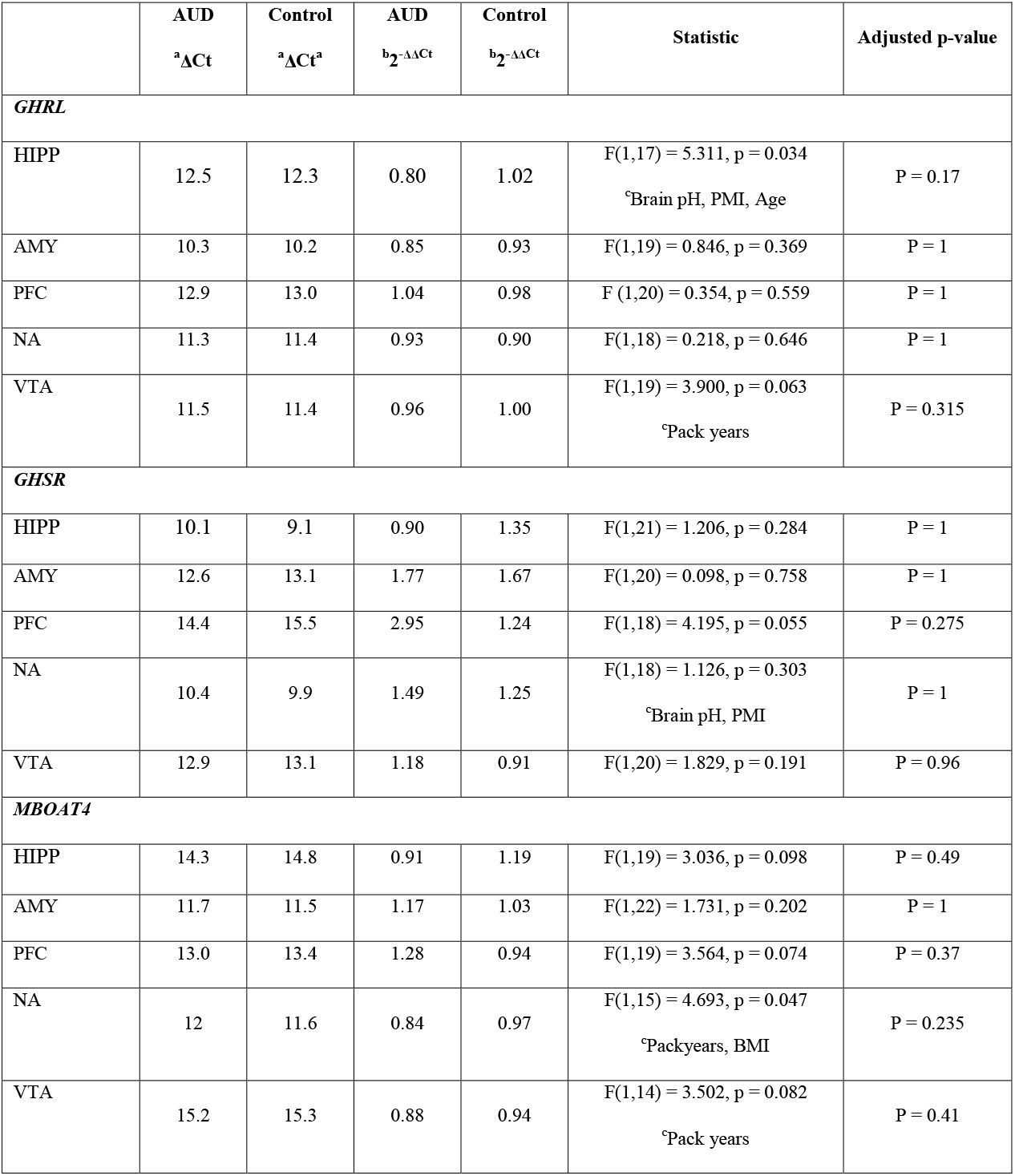
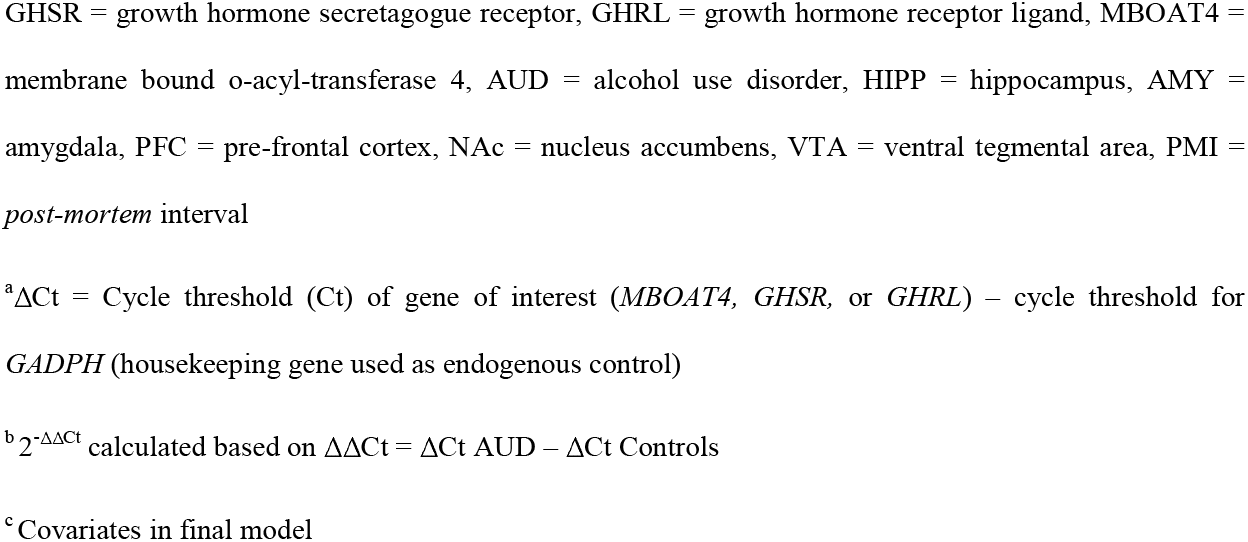
Comparison of *GHRL, GHSR,* and *MBOAT4* expression changes in *Post-mortem* Brain Tissue between AUD and Controls

| | AUD<br><sup>a</sup>ΔCt | Control<br><sup>a</sup>ΔCt<sup>a</sup> | AUD<br><sup>b</sup>$2^{-\Delta\Delta$Ct | Control<br><sup>b</sup>$2^{-\Delta\Delta$Ct | Statistic | Adjusted p-value |
| --- | --- | --- | --- | --- | --- | --- |
| ***GHRL*** |  |  |  |  |  |  |
| HIPP | 12.5 | 12.3 | 0.80 | 1.02 | F(1,17) = 5.311, p = 0.034<br><sup>c</sup>Brain pH, PMI, Age | P = 0.17 |
| AMY | 10.3 | 10.2 | 0.85 | 0.93 | F(1,19) = 0.846, p = 0.369 | P = 1 |
| PFC | 12.9 | 13.0 | 1.04 | 0.98 | F (1,20) = 0.354, p = 0.559 | P = 1 |
| NA | 11.3 | 11.4 | 0.93 | 0.90 | F(1,18) = 0.218, p = 0.646 | P = 1 |
| VTA | 11.5 | 11.4 | 0.96 | 1.00 | F(1,19) = 3.900, p = 0.063<br><sup>c</sup>Pack years | P = 0.315 |
| ***GHSR*** |  |  |  |  |  |  |
| HIPP | 10.1 | 9.1 | 0.90 | 1.35 | F(1,21) = 1.206, p = 0.284 | P = 1 |
| AMY | 12.6 | 13.1 | 1.77 | 1.67 | F(1,20) = 0.098, p = 0.758 | P = 1 |
| PFC | 14.4 | 15.5 | 2.95 | 1.24 | F(1,18) = 4.195, p = 0.055 | P = 0.275 |
| NA | 10.4 | 9.9 | 1.49 | 1.25 | F(1,18) = 1.126, p = 0.303<br><sup>c</sup>Brain pH, PMI | P = 1 |
| VTA | 12.9 | 13.1 | 1.18 | 0.91 | F(1,20) = 1.829, p = 0.191 | P = 0.96 |
| ***MBOAT4*** |  |  |  |  |  |  |
| HIPP | 14.3 | 14.8 | 0.91 | 1.19 | F(1,19) = 3.036, p = 0.098 | P = 0.49 |
| AMY | 11.7 | 11.5 | 1.17 | 1.03 | F(1,22) = 1.731, p = 0.202 | P = 1 |
| PFC | 13.0 | 13.4 | 1.28 | 0.94 | F(1,19) = 3.564, p = 0.074 | P = 0.37 |
| NA | 12 | 11.6 | 0.84 | 0.97 | F(1,15) = 4.693, p = 0.047<br><sup>c</sup>Packyears, BMI | P = 0.235 |
| VTA | 15.2 | 15.3 | 0.88 | 0.94 | F(1,14) = 3.502, p = 0.082<br><sup>c</sup>Pack years | P = 0.41 |

GHRSR = growth hormone secretagogue receptor, GHRL = growth hormone receptor ligand, MBOAT4 = membrane bound o-acyl-transferase 4, AUD = alcohol use disorder, HIPPO = hippocampus, AMY = amygdala, PFC = pre-frontal cortex, NAc = nucleus accumbens, VTA = ventral tegmental area, PMI = *post-mortem* interval
<sup>a</sup>ΔCt = Cycle threshold (Ct) of gene of interest (*MBOAT4*, *GHRSR*, or *GHRL*) – cycle threshold for *GADPH* (housekeeping gene used as endogenous control)
<sup>b</sup> $2^{-\Delta\Delta Ct}$ calculated based on $\Delta\Delta Ct = \Delta Ct \text{ AUD} - \Delta Ct \text{ Controls}$
<sup>c</sup> Covariates in final model

### 2.3 Acyl-Ghrelin Levels are Reduced by Ethanol in Rats, Independent of Ghrelin Receptor Knockout (KO)

We next examined whether a change in ghrelin levels as a result of ethanol administration, as observed in our human laboratory experiments, could be replicated in rodents, and whether this effect would depend on the presence of the ghrelin receptor. We evaluated changes in plasma acyl- and des-acyl-ghrelin levels 15 min following an intraperitoneal (i.p.) injection of ethanol (1.5 g/kg) or saline in wild-type (WT) and ghrelin receptor knockout *(Ghsr* KO) male rats. Using two-way ANOVA, we observed a significant main effect of treatment (ethanol *vs.* saline) on acyl-ghrelin levels [F(1,29) = 6.212, p = 0.019], where levels of acyl-ghrelin in ethanol-treated rats were reduced compared with saline-treated rats. There was no main effect of genotype (WT *vs.* KO) [F(1,29) = 0.2309, p = 0.63] or treatment × genotype interaction [F(1,29) = 0.01, p = 0.92] (**Figure 4A**). Moreover, there was no effect of treatment [F(1,29) = 0.013, p = 0.91], genotype [F(1,29) = 0.3245, p = 0.57], or treatment x genotype interaction [F(1,29) = 0.003, p = 0.95] on plasma des-acyl-ghrelin levels (**Figure 4B**).

### 2.4 Ghrelin Secretion from Gastric Mucosa Cells is Not Altered by Ethanol

Given that we found ethanol reduced acyl-ghrelin secretion in both human and animal experiments, we hypothesized that ethanol may have a direct effect on ghrelin release from the gastric mucosal cells. To test this hypothesis, we evaluated ghrelin release from gastric mucosal cells in the presence of different concentrations of ethanol. We cultured cells in both 5 mM glucose environment and 0 mM glucose environments. A 5 mM glucose condition was used to study the effect of ethanol in settings simulating physiological blood glucose concentrations. A 0 mM glucose condition was used to study the effect of ethanol without any interference of glucose as an energy source, and is known to be associated with higher ghrelin secretion as compared to 5 mM glucose [67]. Because it has been shown to stimulate ghrelin secretion from primary cultures of gastric mucosal cells, norepinrephine (10 μM) was used as a positive control for ghrelin secretion [68]. As observed previously [67], absence of glucose (0 mM *vs.* 5 mM) increased acyl-ghrelin secretion from primary cultrues of gastric mucosal cells. However, ethanol did not change acyl-ghrelin secretion at any of the concentrations tested (**Figure 5A-C**).

**Figure 4:**
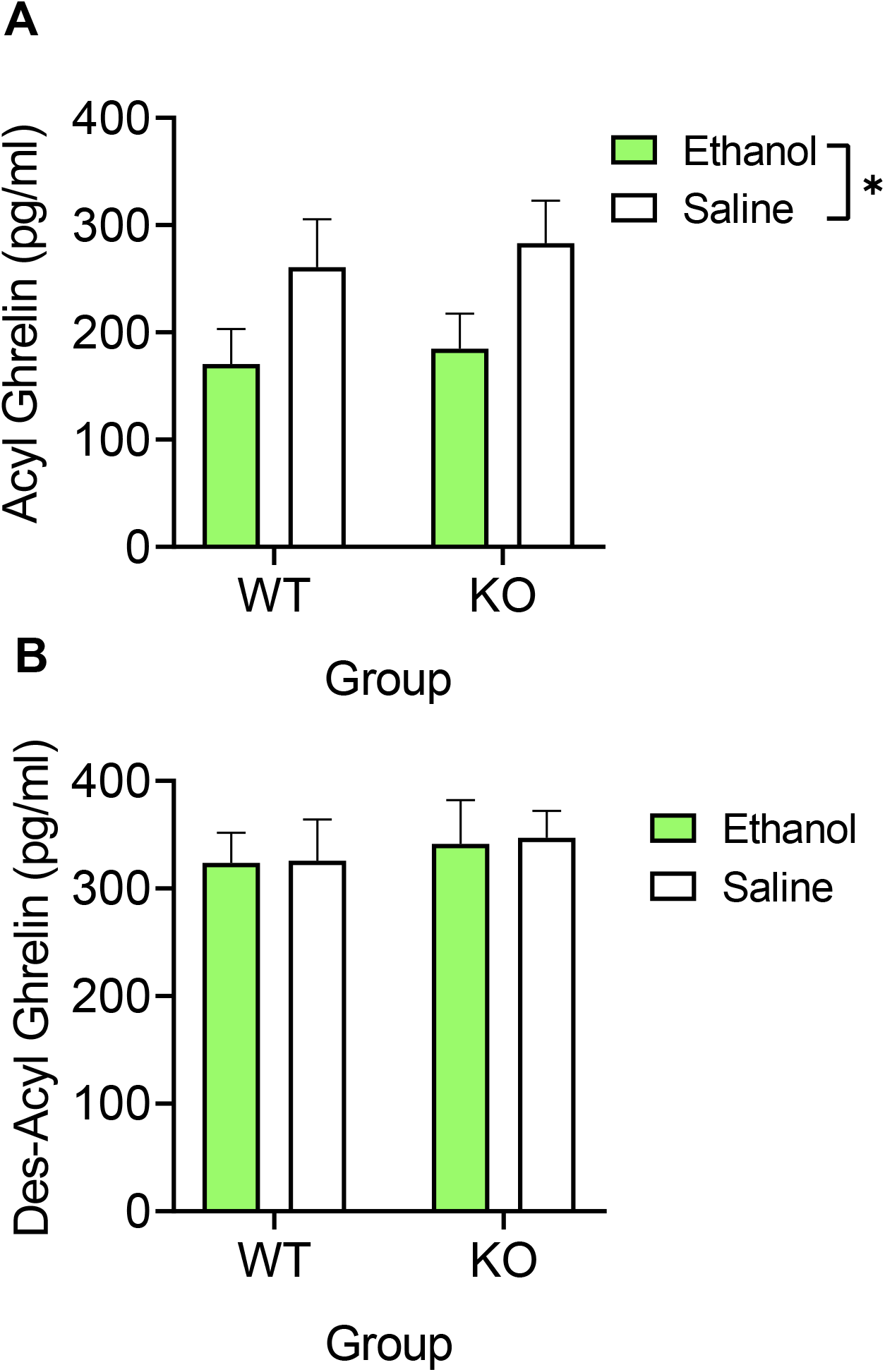
Effect of Ethanol on Peripheral Ghrelin Levels in *Ghsr* KO and WT Rats. **4A:** Effect of ethanol (1.5 g/kg, i.p.) on plasma acyl-ghrelin levels in *Ghsr* KO and WT rats. Data represents change in ghrelin secretion with ethanol treatment *vs.* saline. N = 8-9/group. Treatment effect: p < 0.019, Genotype effect: p = *NS*, and Interaction effect: p = *NS.* **4B:** Effect of ethanol (1.5 g/kg, i.p.) on plasma des-acyl-ghrelin levels in *Ghsr* KO and WT Rats. Data represent change in ghrelin secretion with ethanol *vs.* saline treatment. N = 8-9/group. Treatment effect: p = *NS*, Genotype effect: p = *NS*, and Interaction effect: p = *NS.* Data are presented as Mean ± SEM. p < 0.05 considered significant. *p < 0.05.

**Figure 5:**
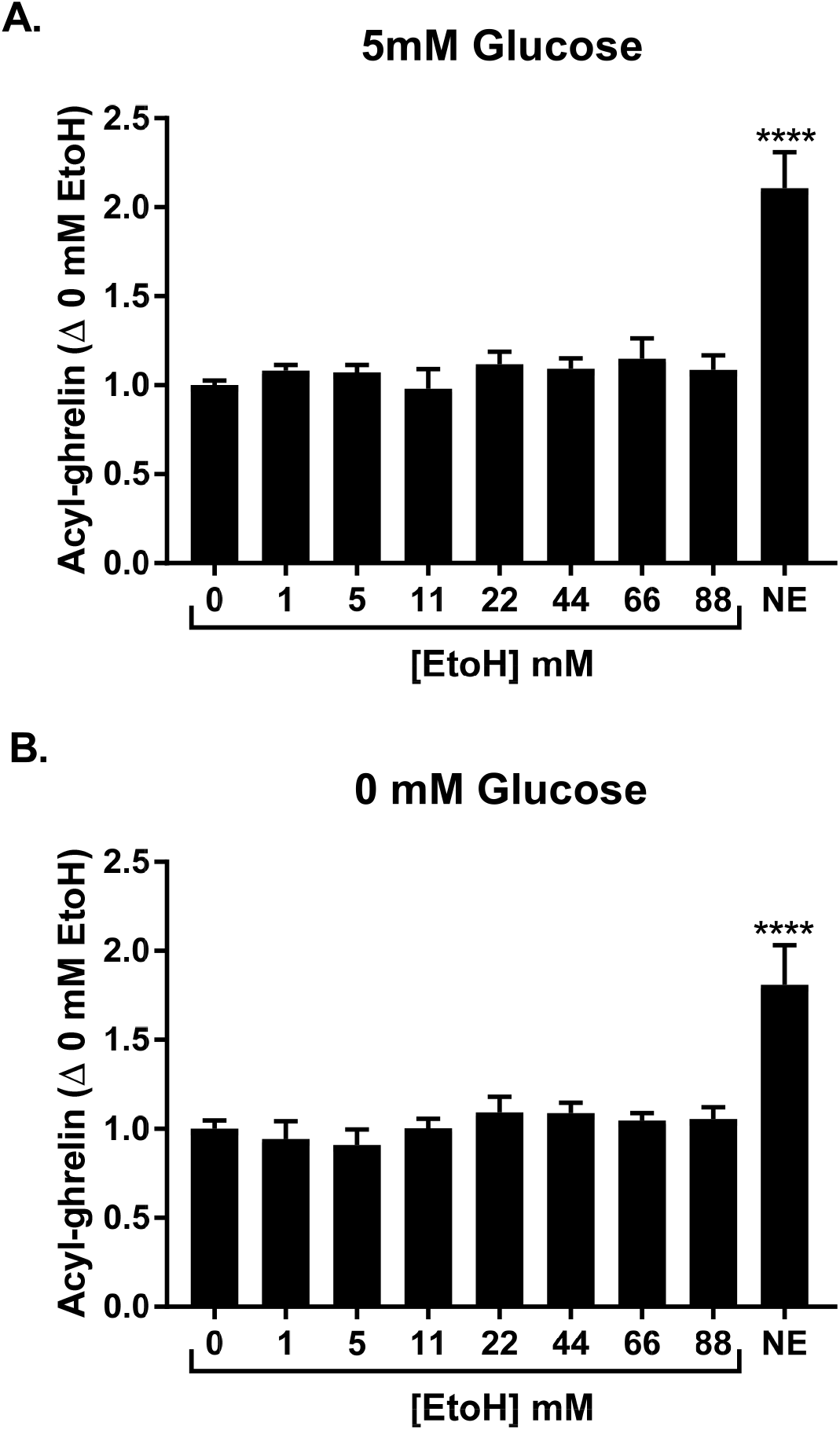
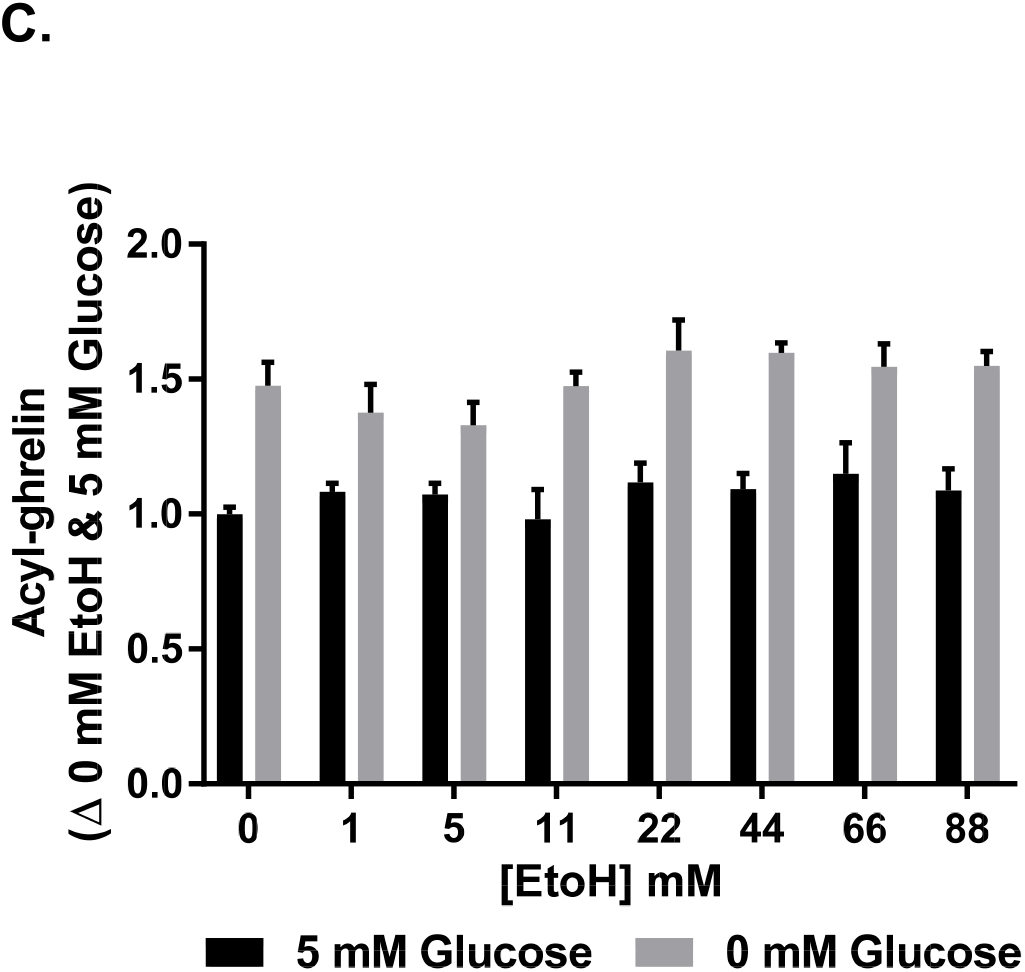
Effect of Ethanol on Ghrelin Secretion from Gastric Mucosal Cells. **5A**: Effect of increasing concentrations of ethanol on acyl-ghrelin secretion in mouse primary gastric mucosal cells incubated with medium containing 5 mM glucose. Data represent change in ghrelin secretion with ethanol (EtoH) treatment compared to untreated control (0 mM EtoH). N = 4 for each condition. ****p < 0.0001, one-way ANOVA followed by Dunnett’s test. **5B**: Effect of increasing concentrations of ethanol on acyl-ghrelin secretion in mouse primary gastric mucosal cells incubated with medium containing 0 mM glucose. Data represents change in ghrelin secretion with EtoH treatment compared to untreated control (0 mM EtoH). N = 4 for each condition. ***p < 0.0001, one-way ANOVA followed by Dunnett’s test. **5C**: Effect of increasing concentrations of ethanol on acyl-ghrelin secretion in mouse primary gastric mucosal cells incubated with medium containing either 5 or 0 mM glucose (for reference only). Data represents change in ghrelin secretion with treatment compared to untreated control (0 mM EtoH in 5 mM glucose). N = 4 for each condition.

### 2.5 Human GOAT (hGOAT) Acylation Activity Is Not Affected by Ethanol

Having shown that ethanol does not affect ghrelin secretion in gastric mucosal cells and that the effects of ethanol on ghrelin levels are independent of the presence or absence of its receptor, we questioned whether ethanol may have direct effects on other components of the ghrelin system, for example, the GOAT enzyme. Therefore, we assayed GOAT activity in increasing concentrations of ethanol. Ethanol was tested at concentrations representing intracellular ethanol levels ranging from sub-intoxicating (1 mM) to grossly intoxicating, lethal (87 mM) doses [69, 70]. We found that ghrelin acylation by GOAT was not dose-dependently inhibited by ethanol over this physiologically relevant concentration range, with less than 20% inhibition observed at the highest concentration tested (**Figure S5**).

### 2.6. Plasma Ghrelin Levels in Rats Are Differentially Affected by Ethanol and Sucrose

Our experiments, thus far, indicated that while ethanol decreases acyl-ghrelin secretion in humans and animals, no direct interaction between ethanol and the ghrelin system (acyl-ghrelin secretion from gastric mucosal cells, GOAT activity, or ghrelin receptor KO) was observed in our aforementioned experiments. We hypothesized that the effect of ethanol to lower peripheral ghrelin concentrations in humans and animals may have occurred indirectly, in proportion to caloric load. We compared the effect of ethanol (1.5 g/kg, ~10 kcal/kg i.p.) with a sucrose solution (2.8 g/kg, ~11.2 kcal/g i.p.) of similar caloric value in male rats, on acyl- and des-acyl-ghrelin in a within-subjects comparison to saline treatment on the previous day.

As for ethanol, we observed a significant main effect of treatment (ethanol *vs.* saline) on acyl-ghrelin levels [F(1, 18) = 7.83, p = 0.02], and a treatment × time interaction effect [F(2,36) = 18.09, p < 0.0001], but no main effect of time. Post-hoc testing revealed a decrease in acyl-ghrelin levels following ethanol treatment, compared to baseline, and in comparison with saline (**Figure 6A, left**). For des-acyl-ghrelin, significant main effects of treatment [F(1,18) = 5.253, p = 0.034] and time [F(1.673, 30.12) = 3.799, p = 0.041], as well as treatment × time interaction effect [F(2, 36) = 3.301, p = 0.048] were observed. Post-hoc testing revealed significant increase in des-acyl-ghrelin levels following saline treatment, but no changes following ethanol treatment (**Figure 6A, right**). There was also a significant main effect of treatment [F(1, 18) = 10.68, p = 0.0043], time [F(2, 36) = 59.03, p < 0.0001], and interaction [F(2, 36) = 50.02, p < 0.0001] on the acyl-to des-acyl-ghrelin ratio (AG:DAG ratio). Post-hoc testing revealed significant reduction in AG:DAG ratio following ethanol treatment, but no changes following saline treatment (**Figure S6**).

**Figure 6:**
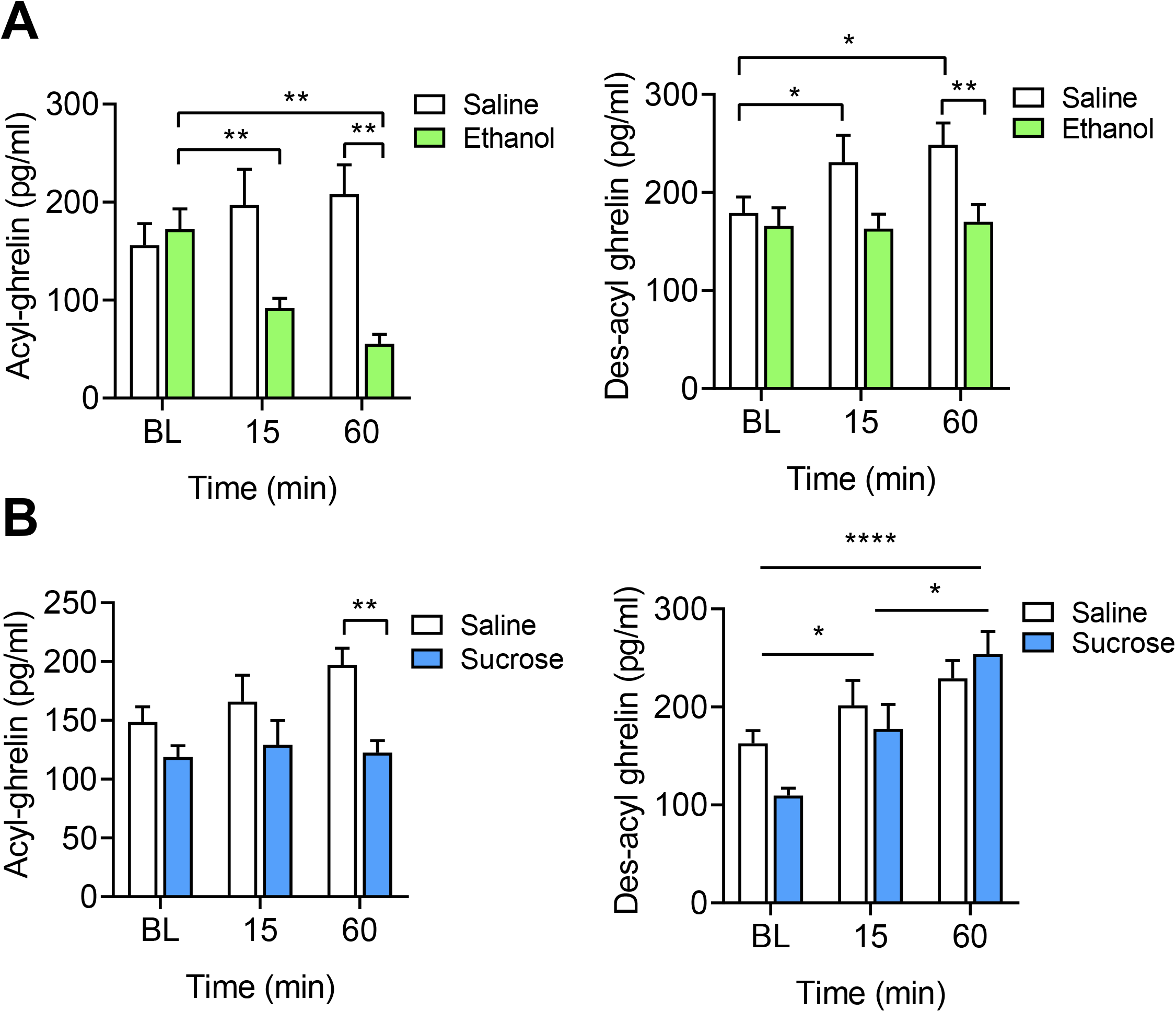
Change in Peripheral Ghrelin as a Result of Alcohol or Sucrose. **6A:** Plasma acyl-ghrelin and des-acyl-ghrelin resulting from alcohol and saline at baseline (BL) and 15 min and 60 min post-injection. *Acyl-ghrelin (left):* Two-way repeated measures (RM) ANOVA: overall treatment (p < 0.05) and interaction (p < 0.0001) effect. Post-hoc testing revealed a decrease in acyl-ghrelin levels at 15 min (p = 0.003) and 60 min (p = 0.004) following ethanol treatment, compared with the pre-treatment baseline. When compared to saline injections, acyl-ghrelin levels were significantly lower in ethanol-treated rats at 60 min post-treatment (p = 0.002). *Des-acyl-ghrelin (right):* Two-way RM ANOVA: overall time (p < 0.05), treatment (p < 0.05), and interaction (p < 0.05). Post-hoc testing revealed significant increases in des-acyl-ghrelin at 15 min (p = 0.047) and 60 min (p = 0.018) following saline treatment, whereas no changes in des-acyl-ghrelin levels were observed following ethanol treatment. Des-acyl-ghrelin levels following saline treatment were significantly higher compared to ethanol treatment at 60 min (p = 0.038). **6B:** Plasma acyl-ghrelin and des-acyl-ghrelin resulting from sucrose and saline treatment at baseline and 15 and 60 min post-inejction. *Acyl-ghrelin (left):* Two-way RM ANOVA: overall treatment (p < 0.05) effect. Post-hoc testing revealed significantly lower acyl-ghrelin levels at 60 min following sucrose treatment, compared to saline treatment (p = 0.002). *Des-acyl-ghrelin (right):* Two-way ANOVA: overall time (p < 0.0001) effect. Post-hoc testing revealed a significant increase in des-acyl-ghrelin at 15 min (p = 0.031) and 60 min (p < 0.001), compared to baseline, and at 60 min (p = 0.0190), compared to 15 min. Post-hoc Sidak’s multiple comparison tests: *p < 0.05, ** p < 0.01, ***p < 0.001, ****p < 0.0001.

As for sucrose, we observed an overall significant main effect of treatment (sucrose *vs.* saline) [F(1, 16) = 7.705, p = 0.01] on acyl-ghrelin levels, but no effect of time or an interaction effect, indicating lower levels of acyl-ghrelin under sucrose treatment, compared to saline, regardless of time (**Figure 6B**, **left**). There was no effect of treatment or treatment × time interaction on des-acyl-ghrelin levels, but a significant effect of time [F(1.839, 29.43) = 20.11, p < 0.0001] was observed, with post-hoc testing indicating an overall increase in des-acyl-ghrelin over time (**Figure 6B, right**). Lastly, we observed a significant main effect of treatment [F(1, 16) = 4.93, p = 0.041], time [F (1.966, 31.45) = 26.38, p < 0.0001], and treatment × time interaction [F (2, 32) = 19.18, p < 0.0001] on the AG:DAG ratio. Post-hoc testing revealed significant reduction in AG:DAG ratio following sucrose treatment, but no, or less robust, changes following saline treatment (**Figure S6**).

## 3. Discussion

The results presented herein demonstrate ethanol administration acutely suppressed both plasma acyl- and total ghrelin levels in humans and rats. While these findings are consistent with previous preliminary work [2, 52–56], they expand our knowledge of the effect of ethanol on ghrelin by providing evidence that this effect occurs independently of the ghrelin receptor, given that ethanol-induced suppression of acyl-ghrelin is observed despite knockout of the ghrelin receptor in rats. Moreover, AUD (associated with long term ethanol use) had no effect on central ghrelin, ghrelin receptor, or GOAT enzyme mRNA expression in select brain regions. Our results also show that the effect of ethanol on acyl-ghrelin secretion does not occur through inhibition of GOAT activity or direct inhibition of gastric mucosal cell secretion of acyl-ghrelin. Moreover, administration of ethanol in comparison to sucrose solution produces distinct effects on peripheral ghrelin, despite similar caloric value. Overall, our data suggest that the mechanism by which ethanol intake suppresses plasma ghrelin is indirect and is not necessarily proportional to caloric load.

We consistently observed a reduction in both acyl-ghrelin and total ghrelin in all four experiments, despite each session employing a different duration, dose of alcohol, and type of alcoholic beverage. Our findings further demonstrate that alcohol administration results in acute suppression of peripheral ghrelin, regardless of the route of administration being oral or intravenous and is therefore irrespective of first-pass metabolism. As such, our work strengthens the suggestion that passage through the stomach is not necessary for acute alcohol-induced ghrelin suppression [56, 57]. We did not observe a statistically significant reduction in acyl-or total ghrelin during the fixed IV administration of alcohol. However, here, we were only able to examine ghrelin levels within 45 min following the start of the session. While other sessions with more timepoints collected revealed that alcohol can significantly decrease acyl-ghrelin within 30 min, the experiment was likely insufficiently powered to detect a change that reached statistical significance within such a short time window. Total ghrelin followed the same trends as acyl-ghrelin for all sessions. Given that we did not have data on des-acyl-ghrelin levels at these timepoints, it is unclear whether the reduction in total ghrelin is mainly reflective of a reduction of acyl-ghrelin, or a reduction of both acyl- and des-acyl-ghrelin in these samples.

Data from *Ghsr* KO and WT rats indicate that intraperitoneal injection of ethanol has no rapid, acute effect on des-acyl-ghrelin and only reduces acyl-ghrelin 15 min post-injection. Moreover, we found that this effect of ethanol on acyl-ghrelin occurred independently of the presence or absence of the ghrelin receptor. Although these results may be due to compensatory mechanisms formed during development, we also show no change in central ghrelin, GOAT, or ghrelin receptor expression as a result of chronic exposure to alcohol in individuals with severe AUD (human *post-mortem* sample). More specifically, *GHRL, GHSR,* and *MBOAT4* mRNA expression levels were not statistically different (after correcting for multiple comparisons) between *post-mortem* brain samples from individuals with AUD and non-AUD controls, indicating that chronic alcohol exposure does not directly affect central ghrelin system expression in these brain regions. However, given that *GHSR* is not highly expressed in the hippocampus, amygdala, PFC, VTA, or NAc, and that our sample size was relatively small, we may not have been able to precisely capture any effect of long-term ethanol use on the ghrelin receptor that was statistically meaningful. It should also be considered that the expression of *GHSR* produces two transcripts, GHSR1a and GHSR1b, the latter of which heterodimerizes with and attenuates GHSR1a [71]. Likewise, expression of *GHRL* can also produce products other than pro-ghrelin [71]. Therefore, these findings are not restricted to ghrelin and its receptor. Furthermore, the significance of central *GHRL* mRNA expression remains to be determined. While some *GHRL* mRNA is found in the brain, and central *GHRL* mRNA translation has been demonstrated in rodents, it remains to be demonstrated whether *GHRL* is centrally translated in humans, with the more significant sites of *GHRL* expression being the stomach and duodenum [72]. Still, our data from rodents and human *post-mortem* samples provide converging, albeit preliminary, evidence that the ghrelin receptor does not play a mediating role in acute alcohol-induced suppression of ghrelin.

*In vitro* assay with the GOAT enzyme, a ghrelin mimetic peptide, and octanoate-CoA revealed no dose-dependent effects of ethanol on GOAT acylation activity. These data indicate that alcohol does not mediate its effects on ghrelin secretion by negatively allosterically modifying or directly inhibiting GOAT acylation activity by interfering with GOAT substrate binding. Moreover, gastric mucosal cell secretion of acyl-ghrelin was unaffected by incubation with ethanol either alone or in the presence of glucose. Cells were also tested in the presence of glucose to eliminate the presence or absence of an energy source as a confounding variable in acyl-ghrelin secretion. As reported previously, absence of glucose increased ghrelin secretion [67, 73], and ethanol had no effect on ghrelin in either condition. We only evaluated secretion of acyl-ghrelin from gastric mucosal cells and did not evaluate any changes in des-acyl-ghrelin, however, from our rodent data, it appears that alcohol primarily suppresses acyl-ghrelin secretion. Taken together, our *in vitro* and *ex vivo* data point toward an indirect mechanism whereby alcohol suppresses peripheral ghrelin levels.

Previous studies have suggested that post-prandial ghrelin suppression occurs in proportion to caloric load and that this may underlie alcohol-induced ghrelin suppression [74–76]. Here, we show that acyl-ghrelin was significantly decreased as a result of ethanol administration both 15-and 60-min following injection compared to baseline, whereas there was no significant change in des-acyl-ghrelin following ethanol treatment between each time point. Following sucrose treatment, there was no change in acyl-ghrelin relative to baseline, but des-acyl-ghrelin was significantly increased relative to baseline at 15- and 60-min. Interestingly, given these effects, ethanol and sucrose had similar effects on the AG:DAG ratio, where a decrease of acyl-ghrelin by ethanol, and an increase of des-acyl-ghrelin by sucrose both decreased the plasma AG:DAG ratio significantly at 15 and 60-min post injection. Both acyl- and des-acyl-ghrelin increased over time among saline treated controls. Relative to saline, acyl-ghrelin was decreased by ethanol, and the increase in des-acyl-ghrelin observed following saline treatment was blunted by ethanol. Sucrose treatment, however, blunted the increase in acyl-ghrelin among saline treated controls and did not change the increase in des-acyl-ghrelin relative to saline. It is possible that the increase in acyl- and des-acyl-ghrelin following saline treatment represents a fasting-induced increase in ghrelin that is differentially affected by ethanol and sucrose. Our data showing that ghrelin is not suppressed in proportion to caloric value alone are supported by studies in humans demonstrating that administration of one type of macronutrient differentially affects ghrelin secretion when compared to a different macronutrient of equivalent caloric value [77, 78]. Moreover, acyl-ghrelin is not dose-dependently decreased by higher doses of IV ethanol (associated with higher caloric value) [57]. Our data also suggest that ghrelin acylation and ghrelin peptide secretion are regulated by separate mechanisms, given the markedly different effects of alcohol and sucrose on these different forms of ghrelin. Acyl-ghrelin plays an important role in relaying meal-related information [79], and it is likely that differences in post-prandial (or post-alcohol) acyl-ghrelin secretion are not simply reflective of calorie content, but represent a more complicated summation of the metabolic effects resulting from a meal or alcohol on energy homeostasis, which can vary according to macronutrients, meal status, and size of meal.

Thus, we report here novel findings on the potential underlying mechanism(s) linking alcohol and the ghrelin system. Our results suggest that an indirect mechanism underlies alcohol-induced suppression of acyl-ghrelin that is unique to alcohol. This finding is not surprising given that alcohol’s inability to be stored causes it to be immediately metabolized in the liver at the expense of other nutrients, competitively inhibiting liver enzymes and depleting the pool of electrons used in oxidation of other nutrients [80]. Therefore, it is likely that the downstream effects of alcohol occur on a faster timescale than other macronutrients, depending on prandial state. We observed reductions in acyl-ghrelin within 15 min following alcohol administration in our rodent experiments, and as early as 30 min in our human experiments, whereas post-prandial reductions in ghrelin reaching statistical significance typically occur between 1-2 h following meal ingestion. Although we did not identify a mechanism by which alcohol suppresses acyl-ghrelin, our data shed further light on the complicated nature of ghrelin-alcohol interplay. Acyl-ghrelin is an important component in the regulation of energy balance by signaling for meal preparation and having long-term protective effects against starvation [26, 27]. To date, models of acyl-ghrelin secretion have identified insulin, glucagon, long chain fatty acids, oxytocin, dopamine, norepinephrine, epinephrine, endocannabinoids, somatostatin, glutamate, and glucose as direct regulators of acyl-ghrelin secretion [73, 81–87], and it is possible that alcohol may decrease acyl-ghrelin indirectly by affecting these targets. Another possibility is that alcohol-induced suppression of acyl-ghrelin results from alcohol’s marked acute inhibition of fatty acid β-oxidation. [80]. Recently, it has been suggested that ghrelin acylation can be supported by β-oxidation of long chain fatty acids to produce medium chain fatty acids able to act as GOAT substrates [83, 86, 88]. Therefore, alcohol-induced inhibition of β-oxidation may be partly responsible for acyl-ghrelin suppression resulting from alcohol intake. Clearly, further studies are needed to identify whether these or other mechanisms, such as indirect suppression through other hormones affected by alcohol, underlie the acute alcohol-ghrelin relationship.

Overall, this set of results contributes to a better understanding of the complex interactions between alcohol and the ghrelin system. Nevertheless, these results should be interpreted in light of the study’s limitations. Data presented from human laboratory experiments are the result of secondary analyses and were, therefore, not designed *a priori* to evaluate the effect of alcohol on peripheral ghrelin (e.g., no control infusion/oral saline administration was performed to compare to alcohol). Additionally, the sample sizes from our human and *post-mortem* experiments are small. Our findings on the central expression of *GHRL, GHSR,* and *MBOAT4* in humans should be further investigated in larger samples. We were unable to evaluate expression of these genes in the hypothalamus, a prominent site of central acyl-ghrelin action that might be more significantly altered by chronic exposure to alcohol. Lastly, it should be noted that our results can currently only be generalized to males, given that the human laboratory experiment samples (all > 70% male) were largely male, and human *post-mortem* samples and animals, used in ethanol administration experiments and for gastric mucosal experiments, were all male.

Our data collectively demonstrate that alcohol affects the ghrelin system by acutely decreasing acyl-ghrelin concentration in the circulation and by blunting fasting-induced increase of plasma des-acyl-ghrelin concentrations, and the mechanism likely occurs independently of the ghrelin receptor, and without direct action on the GOAT enzyme or acyl-ghrelin secretion from gastric mucosal cells. Additionally, ethanol and sucrose in equivalent caloric amounts do not have the same effect on peripheral ghrelin, differentially affecting acyl- and des-acyl-ghrelin relative to baseline and saline-treated controls. Therefore, this study suggests that alcohol acutely suppresses ghrelin without directly interacting with the ghrelin system and not simply according to calorie content of alcohol. While further studies are needed to uncover this mechanism of alcohol-induced ghrelin suppression, our data provide new insight into how these effects occur.

## 4. Methods and Materials

### 4.1 Effects of Alcohol on Peripheral Ghrelin Levels in Humans

To examine the effect of alcohol on endogenous ghrelin levels, we performed separate analyses of four human laboratory experiments conducted by our team at the National Institutes of Health (NIH) Clinical Center in Bethesda, Maryland. These experiments were originally performed as part of three placebo-controlled trials [49, 65, 66] (ClinicalTrials.gov: NCT02039349, NCT01779024, NCT01751386) and included the administration of alcohol to non-treatment-seeking, heavy-drinking individuals, as well as measurement of plasma ghrelin levels. Here, we only included data from the placebo conditions of these experiments. Participants provided informed consent and were compensated for participating in each study. The eligibility criteria of each parent study can be found in the **Supplementary Information** (**S1A, S2A, S3A**) and baseline characteristics of each sample analyzed here can be found in **Table S1**. We analyzed data separately for each of the following experiments: (1) variable dose (priming and self-administration) oral alcohol [65], (2) fixed-dose oral alcohol [66], (3) IV variable dose (self-administration) alcohol [49], and (4) fixed-dose intravenous (IV) alcohol [49]. Detailed descriptions of these studies and their primary outcomes have been previously reported [49, 65, 66]. Descriptions of standardized meals for each study can be found in the supplement (**S1B, S2B, S3B**). An overview of these experiments, including information about times of blood draws, meals, and alcohol administration can be found in **Figure 1** and **Table S2**. Below we provide a brief description of each experiment.

#### 4.1.1 Oral Variable Dose Alcohol Administration

The main aim of the parent study was to test the role of baclofen on alcohol drinking using a randomized, between-subjects, double-blind, placebo-controlled human laboratory design [65]. Here, we analyzed data from the placebo group only (participants randomized to placebo and not baclofen). Participants received their assigned study medication (placebo only in this analysis) for approximately a week before returning to complete the experimental session. Participants were instructed to abstain from alcohol 24 h prior to the experiment (verified by BrAC = 0g/dL) and to take their first medication dose before arriving at the clinic. The experimental session consisted of alcohol cue reactivity followed by alcohol priming and alcohol self-administration (for full details, see: [65]). Briefly, for the alcohol priming, participants were provided with their preferred choice of alcohol, mixer (**S1C, S2C**), and television program. The amount of alcohol in the priming drink was calculated to raise each participant’s BAC to 0.03g/dL [89]. Participants were asked to consume the entire drink within five min. The alcohol self-administration (ASA) session began 40 min after consumption of the priming drink. At the beginning of the ASA session, a tray containing four mini-drinks was offered. Each mini-drink had half the amount of alcohol as the priming drink (BAC increase of 0.015g/dL), and participants were allowed to drink as many of the mini-drinks as they chose, with the knowledge that they would receive $3 for each mini-drink not consumed. An additional tray with 4 mini-drinks was offered again 60 min after the beginning of the ASA session. The total ASA session lasted for 120 min, during which participants were not allowed to exceed a BAC of 0.12 g/dL, and BrAC and vital signs were collected every 30 min. Following completion of the ASA session, participants were escorted to an inpatient unit where they were monitored until BrAC reached 0 g/dL, and they were discharged the next morning.

#### 4.1.2 Oral Fixed-Dose Alcohol Administration

The main aim of the parent study was to test the safety of a ghrelin receptor blocker (PF-5190457), co-administered with alcohol, using a Phase 1b, within-subjects, dose-escalating, single-blind, placebo-controlled human laboratory design [66]. Here, we analyzed data from the placebo condition only. The alcohol administration experiment was held on the third day of an inpatient visit, after taking five doses of the study drug (placebo only in this analysis). A standardized alcoholic beverage (Smirnoff vodka, 40% alcohol by volume; 80% proof) was administered, and participants were instructed to drink the beverage within 15 min. Alcohol was provided as a mixed drink containing the participants’ choice from a list of 7 common mixers (**S1C**). Alcohol administration was designed to bring each participant’s blood alcohol concentration to a target BAC of 0.06 g/dL [89].

#### 4.1.3 Fixed Dose IV Administration of Ethanol and IV-Alcohol Self-Administration

The parent study under which both IV-alcohol experiments were performed was a cross-over, randomized, double-blind, placebo-controlled study testing the effects of exogenous ghrelin administration and consisting of four experimental sessions: two IV ASA (1 ghrelin, 1 placebo) sessions and two brain fMRI sessions (1 ghrelin, 1 placebo). We analyzed data from the placebo sessions only. Participants were admitted to the NIAAA inpatient unit at the NIH Clinical Center on the evening before each experiment day. Before each experiment, an IV catheter was inserted into each arm (one for ghrelin/placebo and one for alcohol infusion/blood sampling). For the placebo conditions (which are the only ones considered in this analysis), saline solution was infused during the entire experiment. IV Alcohol was given as 100% dehydrated ethanol diluted by saline to 6.0% (v/v).

##### 4.1.3.1 IV-ASA Experimental Session

For the IV-ASA experiment, participants were given the opportunity to press a button to receive IV-alcohol infusions using a Computerized Alcohol Infusion System (CAIS) during a 120-min session. A progressive-ratio schedule for self-administration was applied, which required the participants to press the button an increasing number of times to receive the subsequent alcohol infusion. Incremental infusion rates were calculated individually to raise each participants BAC to 0.0075mg/dL within two min [89].

BrAC measurements were taken every 15 min throughout the procedure and entered in CAIS software for model-based algorithm adjustments and BAC prediction. Participants were not allowed to exceed a BrAC of 0.12 g/dL during the ASA session. For additional detail, please see: [49, 90, 91]

##### 4.1.3.2 Fixed Dose IV-Administration Experiment

The fixed-dose IV-alcohol administration was conducted as part of a brain fMRI experiment in which subjects completed an alcohol-food incentive delay (AFID) task that exposed participants to food, alcohol, and neutral symbols. After the loading dose, participants completed an initial AFID task without exposure to alcohol. Participants then received continuous IV alcohol infusion while they repeated the resting state scan and AFID task again. The alcohol infusion was given to raise each participant’s BrAC linearly to 0.08 g/dL[89], within 20 min, and clamp the BrAC at this target value until the end of the experiment. The total duration of the IV alcohol infusion was 35 min. For additional detail, please see [49, 90, 91].

#### 4.1.4 Clinical Blood Collection, Processing, and Assay of Ghrelin Levels

For each experiment listed above, blood was collected at multiple time points throughout each experimental session to allow for repeated measures of plasma acyl-and total-ghrelin levels (**Figure 1, Table S2**). The full technical details for blood collection, plasma extraction, and acyl and total ghrelin assays for each study can be found in the Supplement (**S1D, S2D, S3C**).

### 4.2 *GHSR, GHRL,* and *MBOAT4* Gene Expression in Human *Post-mortem* Brain Tissue

Expression levels of the ghrelin receptor gene *(GHSR),* ghrelin gene *(GHRL),* and GOAT gene *(MBOAT4)* were analyzed in *post-mortem* brain samples from male subjects diagnosed with severe alcohol use disorder (AUD) (DSM-5) and controls who did not have a diagnosis of AUD. Human *postmortem* brain tissue was obtained from the New South Wales Tissue Resource Centre (NSWBTRC) at the University of Sydney, Australia [92]. *GHSR, GHRL,* and *MBOAT4* RNA extraction, reverse transcription, and qPCR analysis were performed using procedures previously reported [93]. mRNA was extracted from the five available brain regions, including amygdala, hippocampus, VTA, NAc, and PFC (superior frontal Brodmann areas 8 and 9). Full technical details can be found in the **Supplementary Information (S4)**.

### 4.3 Effects of Ethanol Administration on Peripheral Ghrelin Levels in *Ghsr* KO and WT Rats

Male WT and *Ghsr* KO (Wistar background) rats weighing 350 – 850g were obtained from the Transgenic Breeding Facility at the National Institutes on Drug Abuse (NIDA) Intramural Research Program (IRP) (Baltimore, MD, USA). The development and characterization of the *Ghsr* KO rat has been previously described [94]. Animals were single-housed and maintained in temperature-controlled facilities on a 12 h/12 h light cycle with standard chow and water available *ad libitum.* Rats from both genotype groups randomly received either ethanol (20% w/v) and saline: (1) *Ghsr* KO – Alcohol (n = 8), (2) *Ghsr* KO – Saline (n = 9), (3) WT – Alcohol (n = 9), and (4) WT – Saline (n = 8). Rats were given an intraperitoneal (i.p.) injection of either (1.5 g/kg, 7.5 ml/kg) or saline (1 ml/kg) 15 min before collection of trunk blood into EDTA coated tubes containing inhibitors appropriate for acyl-ghrelin/des-acyl-ghrelin measurement. Full technical details of processing and assay can be found in the **Supplementary Information (S5).**

### 4.4 The Effect of Ethanol on Ghrelin Secretion from Gastric Mucosal Cells

Gastric mucosal cells were isolated and established from 8-12 wk-old male C57BL/6N mice as reported previously [68, 73], and then supplemented with sodium octanoate-bovine serum albumin (BSA) before they were treated with medium containing different ethanol concentrations. After incubation, mediums were assayed for acyl-ghrelin by ELISA. Full technical details can be found in the **Supplementary Information (S6)**.

### 4.5 Effect of Ethanol on Human GOAT (hGOAT) Ghrelin Acylation Activity

Assays were performed with 70 μg membrane protein from Sf9 cells expressing hGOAT, as determined by Bradford assay. Each ethanol concentration was tested by adding an ethanol stock to a mix of HEPES, membrane protein, and MAFP before initiating reactions with octanoyl-coA and GSSFLC_AcDan_ peptide. Reactions were stopped after 1hr with acetic acid, and the medium was analyzed using reverse-phase HPLC, as described previously [95, 96]. GOAT acylation activity was determined by substrate and product peak integration in the presence of either ethanol or water (vehicle). Percent activity for each reaction was calculated using equations 1 and 2 [97]. Full technical details can be found in the **Supplementary Information (S7).**

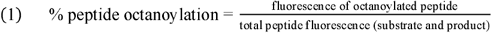

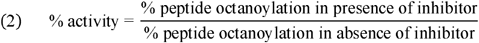

### 4.6 Effects of Ethanol Administration on Peripheral Ghrelin Levels in Rats

Male Wistar rats 10-12 months old (400-700 g) were obtained from Charles River Laboratory (Wilmington, MA). Animals were single-housed and maintained in temperature-controlled facilities on a 12 h/12 h light cycle with standard chow and water available *ad libitum.* Prior to the experiment, rats were habituated to i.p. injections for three days by performing daily saline injections. On the day before the experiment, rats were also habituated to the testing room for 1 h. The following day, baseline measures for each rat were collected via tail bleed 6-7 h into the light cycle, at 15 and 60 min, following saline injection (1 ml/kg, i.p.). Rats were returned to their home cages in between injection and blood draws. Food and water remained accessible to the rats in the home cages. The following day, rats were divided into two groups and received either 20% w/v ethanol (1.5 g/kg, 7.5 ml/kg, i.p.) or 35% w/v sucrose (2.8 g/kg, 8 ml/kg, i.p.). Tail blood was again drawn at the 0, 15, and 60 min timepoints, and processed as described in Experiment 3.3.(**S5**).

### 4.7 Statistics

*Human laboratory experiments:* outliers (defined as outside of ± 1.5(IQR) per hormone per timepoint) were removed. Data were analyzed using Linear Mixed Effects Models in SPSS 25 (IBM Corporation, Armonk, NY) and were evaluated for random effect of subject, main effect of time point, and covariates (age, gender, BMI, and race) on acyl-or total ghrelin. Random effects were described with a scaled identity covariance structure. Covariates that were not significant in the initial run of each model were removed from the final model. Post-hoc analyses were performed using pairwise comparisons of group means at each timepoint within an experiment, and Bonferroni correction was used to conservatively control for multiple comparisons. *Human post-mortem experiment:* Human *post-mortem* data were analyzed using Linear Mixed Effects Models in SPSS 25 and were evaluated for random effect of subject, fixed effect of group (AUD, Non-AUD), and covariates *(Post-mortem* interval (PMI), age, brain weight, brain pH, BMI, and cigarette pack years) on fold change (2^-ΔΔCt^) mRNA expression in each brain region. Covariates that were not significant in the initial run of each model were removed from the final model. A variance components covariance structure was used to describe random effects. To conservatively control for multiple comparisons, Bonferroni correction was applied to correct for the number of brain regions tested. *Ethanol experiments with KO and WT rats:* A two-way ANOVA was used to evaluate genotype (KO *vs.* WT), treatment (ethanol *vs.* saline), and genotype x treatment interaction main effects among saline and ethanol treated *Ghsr* KO and WT rats in GraphPad Prism. Tukey’s multiple comparison test was used to evaluate differences in group means between all genotype and treatment groups. *Gastric mucosal cell experiments:* For ghrelin secretion studies in gastric mucosal cells, a one-way ANOVA followed by Dunnet’s test was used to analyze the effect of different concentrations of ethanol on acyl-ghrelin secretion in Graph Pad Prism. *Ethanol and sucrose experiment with rats:* Lastly, two-way repeated measures ANOVA was used to evaluate treatment (ethanol *vs.* saline and sucrose *vs.* saline), time (0, 15, and 60 min), and treatment x time interaction main effects on acyl-ghrelin and des-acyl-ghrelin in rats using GraphPad Prism. Sidak’s multiple comparison test was used for post-hoc analyses. For all analyses, significance was set at p < 0.05.

### 4.8 Approvals

Human laboratory experiments were approved by the NIH Addictions Institutional Review Board, registered at ClinicalTrials.gov (NCT02039349, NCT01779024, NCT01751386), reviewed by the FDA if applicable under Investigational New Drug (IND) applications, and conducted in accordance with Declaration of Helsinki principles. All participants provided written informed consent before any protocol-specific research procedure took place. The human *post-mortem* brain project was approved by the NIAAA Scientific Advisory Board and the NIH Office of Human Subjects Research Protections and was exempt from review by the NIH Institutional Review Board. Animal studies performed at the NIH IRP adhered to the National Research Council *Guide for the Care and Use of Laboratory Animals* and were approved by the Institutional Animal Care and Use Committee of the NIDA IRP. All animal procedures and use of mice at UT Southwestern Medical Center (UTSW) were approved by the Institutional Animal Care and Use Committee of UTSW.

## Supporting information

NIH Cover Sheet

## Author Contributions

*Overall study basis, rationale and concept:* LL

*Design of the experiments:* 1) current analyses from the human laboratory experiments (MF, LL), 2) *in vivo* rodent experiments (MF, AGF, LJZ, RCNM, BJT, GFK, LFV, LL); 3) *ex vivo* experiments in gastric mucosal cells (JMZ); 4) *in vitro* assays of GOAT enzyme activity (JEM, MR, JLH).

*Acquisition and management of data:* 1) human laboratory experiments (MF, SLD, MRL, FA, LL), 2) *post-mortem* experiments (MF, SLD, HS, MRL), 3) *in vivo* rodent experiments (AGF, LJZ, RCNM, BJT, LFV); 4) *ex vivo* experiments in gastric mucosal cells (BKM, JMZ); 5) *in vitro* assays of GOAT enzyme activity (JEM); 6) human ghrelin assays (FA); and 7) rodent ghrelin assay (ZBY).

*Analysis and interpretation of data:* 1) human laboratory experiments (MF, SLD, LL), 2) *post-mortem* experiments (MF, SLD, LL), 3) *in vivo* rodent experiments (MF, SLD, AGF, LJZ, RCNM, BJT, GFK, LFV, LL); 4) *ex vivo* experiments in gastric mucosal cells (BKM, MR, JMZ); 5) *in vitro* assays of GOAT enzyme activity (JEM, MR, JLH); 6) human ghrelin assays (MF, SLD, FA, LL); and 7) rodent ghrelin assay (MF, SLD, AGF, LJZ, RCNM, BJT, GFK, LFV, LL)

*Clinical and safety monitoring for the human studies:* MRL, LL

*Provided funding:* FA, JLH, JMZ, GFK, LL

*Drafting the manuscript:* SLD

*Assisted with drafting the manuscript:* MF, DMH, LL

All authors have critically reviewed the manuscript for important intellectual content and approved the final version of the manuscript.

## Acknowledgements

We thank the clinical and research staff involved in patient care, data collection/analysis, and technical support in the joint NIDA/NIAAA Clinical Psychoneuroendocrinology and Neuropsychopharmacology Section, in the NIAAA clinical program of the Division of Intramural Clinical and Biological Research (DICBR) (in particular the NIAAA Office of the Clinical Director and the NIAA Clinical Core Laboratory), at the NIH Clinical Center (Departments of Nursing, Nutrition, and Pharmacy), and in the Clinical Pharmacokinetics Research Laboratory at the University of Rhode Island.

We would also like to thank Dr. Melanie Schwandt (Office of the Clinical Director, NIAAA) for data management. We would like to thank Dr. Vijay Ramchandani (Section on Human Psychopharmacology, NIAAA DICBR) and Dr. Reza Momenan (Clinical NeuroImaging Research Core, NIAAA DICBR) for their support in the execution of the parent studies from which these analyses stemmed. The authors would also like to express their gratitude to the participants who took part in these studies. We would like to thank the members of the Neurobiology of Addiction Section, the Transgenic Breeding Facility and the Genetic Engineering and Viral Vector Core in the Intramural Research Program at NIDA/NIH involved in animal care and technical support. Finally, the authors would like to thank Ms. Donna Sheedy and Dr. Jillian Kril from the New South Wales Tissue Resource Centre (NSWBTRC) at the University of Sydney, Australia, for providing the human *post-mortem* brain tissue for this project. The content of this article is solely the responsibility of the authors and does not necessarily represent the official views of the National Institutes of Health.

