## Supplemental Material for "Pursuing the Mechanisms Underlying Alcohol-Induced Changes in the Ghrelin System: New Insights from Preclinical and Clinical Investigations"

**Supplementary Information**

**Appendix S1. Oral Alcohol Priming and Alcohol Self-Administration (ASA) Experiment**

**S1.A. Eligibility Criteria**

***Inclusion Criteria:***

- Male or female between 21 and 65 years old
- Current diagnosis of alcohol dependence (AD) according to the Structured Clinical Interview for DSM-IV-TR Axis I Disorders (SCID)
- A score of ≥ 40 on the trait version of Spielberger State Trait Anxiety Inventory (STAI)
- No or mild alcohol withdrawal symptoms, defined as Clinical Institute Withdrawal Assessment for Alcohol-Revised (CIWA-Ar) score ≤ 8
- Good health as confirmed by medical history, physical examination, ECG, blood/urine lab tests
- Females only: postmenopausal for at least one year, surgically sterile, or practicing an effective method of birth control before entry and throughout the study; reliable methods of birth control include: oral contraceptives or Norplant®; barrier methods such as diaphragms with contraceptive jelly, cervical caps with contraceptive jelly, condoms with contraceptive foam, or intrauterine devices; a partner with a vasectomy; abstinence from intercourse

***Exclusion Criteria:***

- Current diagnosis of substance dependence (other than alcohol and nicotine) based on SCID
- Lifetime diagnosis of schizophrenia, bipolar disorder, or other psychoses; lifetime history of attempted suicide; a diagnosis of Major Depressive Disorder (MDD) within the past 6 months
- History of epilepsy or alcohol-related seizures
- Express interest in treatment for alcohol use disorders and/or anxiety at any time during the study
- Positive urine test for drugs of abuse at any time during the study
- Females only: breast-feeding and/or positive urine pregnancy test at any time during the study
- Poor venous access based on physical exam and medical history
- History of adverse reaction or hypersensitivity to baclofen
- Medical contraindications for baclofen use
- Clinically significant ECG abnormalities, uncontrolled hypertension, Creatinine > 2 mg/dL, and/or clinically significant liver problems (*i.e.*, liver cirrhosis, AST or ALT > 5x the upper normal limit, Hepatitis B or C)
- Current use of psychotropic medications that cannot be discontinued and may have an effect on alcohol consumption or may interact with baclofen, including: naltrexone, acamprosate, alcohol dehydrogenase inhibitors, topiramate, gabapentin, ondansetron, benzodiazepines, β-blockers, H_2_-blockers, and α_1_-blockers

**S1.B. Standardized Meal Options**

**Breakfast** (standard calorie: 400 kcal):

- Option A: 1 cornflakes box, 1 banana, 1 apple sauce, 1 orange juice (4 fl oz), 1 non-fat milk (8 fl oz), 2 sugar packets. (13% protein, 2% fat, and 85% carb)
- Option B: 1 cottage cheese, 2 hardboiled eggs, 2 slices American cheese, 2 slices deli turkey, 1 bottled water. (43% protein, 47% fat, and 10% carb)

**Lunch** (standard calorie: 400 kcal):

- Option A: ½ sandwich (whole wheat bread, deli turkey, and American cheese), 1 potato chips bag, 1 bottled water. (19% protein, 45% fat, and 36% carb)
- Option B: ½ sandwich (multigrain bread, peanut butter, and strawberry jelly), 1 banana, 1 bottled water. (9% protein, 26% fat, and 65% carb)
- Option C: salad (green leaf lettuce, tomato, and cucumber), 1 packet of Italian salad dressing, 1 plain yogurt, 1 apple, 1 bottled water. (17% protein, 29% fat, and 54% carb)

**Dinner** (not standardized): Each participant picked one entrée and one dessert; all participants got all of the side dishes.

- Entrée options: spaghetti with meat sauce, roast turkey with mashed potatoes and gravy, chicken Caesar salad with croutons.
- Dessert options: chocolate cake, apple pie, angel food cake, blueberry yogurt
- Side dishes: macaroni and cheese, steamed broccoli, steamed carrots, dinner roll, and 1 bottled water

**S1.C. Alcohol and Mixer Options**

Participants could choose from 7 options of mixers (see below). Data regarding the preferred brand and other features (e.g., pulp or no pulp) were also collected. Alcohol provided during experimental session was selected based on participant’s personal preference determined at the baseline study visit. Participants could choose any type and brand of alcohol. A participant’s preferred mixer was only provided to participant where requested (i.e., participant’s preferred alcoholic beverage was liquor with cranberry juice, etc.)

- Option 1: Cranberry juice

(If the participant did not have a preference, Ocean Spray Cranberry Juice Cocktail was provided.)

- Option 2: Fruit punch

(If the participant did not have a preference, Hawaiian Punch was provided.)

- Option 3: Lemonade

(If the participant did not have a preference, Newman’s Own Virgin Lemonade was provided.)

- Option 4: Orange juice

(If the participant did not have a preference, Tropicana Pure Premium Orange Juice - Original with No Pulp was provided.)

- Option 5: Pineapple juice

(If the participant did not have a preference, Dole Pineapple Juice was provided.)

- Option 6: Tonic water

(If the participant did not have a preference, Schweppes Tonic Water was provided.)

- Option 7: Soda

(If the participant did not have a preference, Shasta Cola was provided.)

**S1.D. Blood Collection, Processing, and Measurement of Acyl- and Total Ghrelin**

For blood collection, a saline lock intravenous catheter was inserted into the antecubital fossa of the non-dominant arm for multiple blood draws. At each time-point, blood was collected into a K_2_EDTA tube (BD Vacutainer®) and centrifuged within 30 min post-collection (relative centrifugal force: 1700×g, temperature: 4°C, centrifugation time:15 min). Tubes used for the collection of plasma for acyl-ghrelin measurement were pre-treated with the following inhibitors: 4-(2-aminoethyl)benzene sulfonyl fluoride hydrochloride (Pefabloc®; Roche Diagnostics GmbH, Germany), dipeptidyl peptidase IV inhibitor (EMD Millipore Corp., Billerica, MA, Cat.# DPP4-010), and protease inhibitor cocktail (Sigma-Aldrich Inc., Saint Louis, MO – Cat. #P8340). This was done in accordance with assay manufacturer recommendations. After centrifugation, plasma was aliquoted into microtubes and immediately stored at -80°C until assay. Microtubes to be used for total-ghrelin measurement were treated with 5% (v/v) HCl before freezing, according to assay manufacturer recommendations. Acyl-ghrelin was measured using a Millipore Human Metabolic Hormone Magnetic Bead Panel MILLIPLEX® MAP kit (EMD Millipore Corp., Billerica, MA – Cat. #HMHEMAG-34K). The assay was performed on fluorescence-coded magnetic beads coated with capture antibodies specific for each marker (GIP, active amylin, active GLP-1, insulin, leptin, PP, and PYY were assayed simultaneously with acyl-ghrelin) and performed according to manufacturer instructions. A MAGPIX® instrument (Luminex Corp., Austin, TX) allowed for simultaneous detection of all antibodies. Milliplex data were pre-processed and analyzed in the MILLIPLEX® Analyst Software (V. 3.5, EMD Millipore Corp., Billerica, MA) to calculate the concentration of each hormone (pg/ml).Total ghrelin was measured using Human (Total) Ghrelin ELISA kits (EMD Millipore Corp., Billerica, MA, Cat.#EZGRT-89K) according to manufacturer instructions with samples measured in duplicate. The optical density of each well was determined using the GloMax®-Multi Detection System (Promega Corp., Madison, WI – Part#TM297) and a regression model was applied to calculate the concentration in MS Excel (Microsoft, Redmond, WA).

**Appendix S2. Fixed Oral Alcohol Administration Experiment**

**S2.A. Eligibility Criteria**

Inclusion criteria:

- Male or female between 21 and 65 years old
- Weekly alcohol consumption of > 15 and > 20 standard drinks for women and men, respectively
- No or mild alcohol withdrawal symptoms, defined as a Clinical Institute Withdrawal Assessment of Alcohol Scale, Revised (CIWA-Ar) score of ≤ 8
- Good health as confirmed by medical history, physical examination, electrocardiogram (ECG), blood/urine lab tests
- Female participants must be of non-childbearing potential, as defined by at least one of the following criteria:
  - 21- to 65-year-old females who have a documented hysterectomy and/or bilateral oophorectomy
  - 45- to 65-year-old females who are postmenopausal, defined as follows:
    - 45- to 55-year-old females who satisfy all the following three criteria:
      1) Amenorrhea, defined as absence of menstruation for the past 12 months
      2) Negative urine pregnancy test
      3) Serum follicle-stimulating hormone (FSH) concentration within the laboratory’s reference range for postmenopausal females
    - 56- to 65-year-old females with amenorrhea, defined as absence of menstruation for the past 12 months
- Male participants must agree to use one of the following contraception methods from the first dose of study drug until 28 days after the last dosing:
  - Abstinence
  - Condom AND one of the following:
    - Vasectomy for more than 6 months
    - Female partner who meets one of the following:
      1) Is postmenopausal
      2) Has had a tubal ligation, hysterectomy, or bilateral oophorectomy
      3) Uses one of the following contraception methods: copper or hormonal intrauterine device (IUD), spermicidal foam/gel/film/cream/suppository, diaphragm with spermicide, oral contraceptive, injectable progesterone, subdermal implant

Exclusion criteria:

- Expressing interest in receiving treatment for alcohol problems at any time during the study
- Positive urine test for drugs of abuse at any time during the study
- Current diagnosis of substance dependence (other than alcohol and nicotine) based on structured clinical interview for DSM-IV
- lifetime history of attempted suicide or self-injury; lifetime diagnosis of schizophrenia, bipolar disorder, or other psychoses; current diagnosis of clinically significant major depressive disorder and/or anxiety disorder based on structured clinical interview for DSM-IV
- Clinically significant medical problems (e.g., unstable hypertension, clinically significant ECG abnormalities, creatinine ≥ 2 mg/dL, liver cirrhosis, aspartate aminotransferase or alanine transaminase > 3× the upper normal limit, hemoglobin < 10.5 g/dl
- Patients who have a diagnosis of diabetes and/or are treated with any drug that has glucose lowering properties, such as sulfonylurea, insulin, metformin, thiazolidinediones (TZD), dipeptidyl peptidase-4 inhibitors, or glucagon-like peptide-1 (GLP-1) agonists
- Heart rate > 100 beats per minute at screening on two separate measurements
- BMI ≥ 35 kg/m^2^ or BMI ≤ 18.5 kg/m^2^ or anorexia
- History of epilepsy or alcohol-related seizures
- History of alcohol-induced flushing
- Medications use:
  - The following medications are exclusionary if taken within 2 weeks prior to the study drug administration: naltrexone, acamprosate, alcohol dehydrogenase inhibitors, topiramate, gabapentin, ondansetron, benzodiazepines, β-blockers, H_2_-blockers, α_1_-blockers, baclofen, barbiturates, hormone replacement therapy, medications and dietary/herbal supplements (e.g., St. John's wort) that interact with cytochrome P450 3A4.
  - Participants that are required to take the following P-glycoprotein inhibitors and inducers are excluded unless they stop taking the inhibitors for 2 weeks and inducers for 6 weeks prior to the study drug administration:
    - Inhibitors: amiodarone, azithromycin, captopril, carvedilol, clarithromycin, conivaptan, cyclosporine, diltiazem, dronedarone, erythromycin, felodipine, itraconazole, ketoconazole, lopinavir, ritonavir, quercetin, quinidine, ranolazine, verapamil
    - Inducers: avasimibe, carbamazepine, phenytoin, rifampin, St John’s wort, tipranavir/ritonavir

**S1.B. Standardized Meal Options**

Participants could choose from 6 options of standardized meals (see below) and were instructed to eat everything served. All meals were individually adjusted for each participant’s calorie needs calculated using the Mifflin-St. Jeor equation [1] with an activity factor of 1.5. All meal options had the same macronutrient distribution of ~20% protein, ~30% fat, ~50% carbohydrate.

For timing of the standardized meals, see **Figure 1** (main manuscript).

*Standardized breakfast:*

- Option 1: Cheerios with milk + Toast with butter + Cottage cheese + Pineapple
- Option 2: Pancakes with butter and syrup + Turkey sausage + Peach yogurt
- Option 3: Scrambled egg + Blueberry muffin + Strawberry yogurt + Peaches
- Option 4: French toast with syrup + Scrambled eggs with cheddar cheese
- Option 5: Pancakes with butter and syrup + Milk + Scrambled eggs
- Option 6: Omelet with swiss cheese, spinach, mushrooms, and onions + Whole-wheat toast with butter and strawberry jelly

*Standardized lunch:*

- Option 1: Roast beef and provolone + Sandwich with mayo, lettuce, and tomato on multigrain bread + Potato chips + Grapes
- Option 2: Grilled chicken and provolone wrap with mayo, lettuce, and tomato + Apple + Pretzels
- Option 3: Grilled chicken Caesar salad + Dinner roll + Applesauce + Chocolate chip cookie
- Option 4: Turkey and swiss sandwich with mayo, mustard, lettuce, and tomato on whole-wheat bread + Baby carrots with ranch dressing + Pretzels+ Tropical fruit
- Option 5: Vegetable quesadilla + Black beans + Rice + Salsa + Orange
- Option 6: Turkey and provolone sandwich with mayo and mustard + Potato chips + Grapes

*Standardized dinner:*

- Option 1: Turkey + Mashed potatoes and gravy + Broccoli + Dinner roll with butter + Chocolate chip cookie
- Option 2: Grilled chicken + Macaroni and cheese + Broccoli + Grapes + Ginger ale
- Option 3: Beef tender roast + Baked potato with butter and sour cream + Green beans + Angel food cake + Peaches
- Option 4: Hamburger + French fries with ketchup
- Option 5: Baked tilapia + Wild rice mix + Sautéed spinach + Dinner roll with butter + Chocolate pudding + Oreos
- Option 6: Spaghetti with marinara, turkey meatballs, and parmesan cheese + Dinner roll + Side salad with Italian dressing

**S2.C. Alcohol and Mixer Options**

Participants could choose from 7 options of mixers presented in S1.C. Smirnoff vodka (40% alcohol by volume) was always used for the alcohol part of the drink.

**S2.D. Blood Collection, Processing, and Assay**

Blood was drawn from the peripheral vein of the arm with a 20 ga IV via a 3-way stopcock. All blood processing, plasma extraction, storage and assay for acyl and total-ghrelin were done following the same procedures listed above in **S1.D.**

**Appendix S3. Fixed Dose IV Alcohol Administration and IV ASA Experiments**

**S3A. Eligibility Criteria**

***Inclusion Criteria:***

- Male or female between 21 and 65 years old
- Weekly alcohol consumption of >15 and >20 standard drinks for women and men, respectively
- No or mild alcohol withdrawal symptoms, defined as a Clinical Institute Withdrawal Assessment of Alcohol Scale, Revised (CIWA-Ar) score of ≤ 8
- Good health as confirmed by medical history, physical examination, electrocardiogram (ECG), blood/urine lab tests
- Willing to have two intravenous (IV) lines during the experimental sessions
- Psycho-social stability (*e.g*., fixed address, reliable person to contact in case of emergency)
- Sexually active females of childbearing potential must agree to use an effective method of birth control during the study. Adequate methods include: oral contraceptive pills plus a barrier method, spermicide plus a barrier method (*e.g*., condom, diaphragm, cervical cap), an intrauterine device (IUD) approved by the Food and Drug Administration (FDA), surgically sterile male partner(s), exclusively female partner(s)

***Exclusion Criteria:***

- Expressing interest in receiving treatment for alcohol problems at any time during the study
- Alcohol-naïve, current alcohol abstainer, or no prior experience of drinking ≥ 5 standard drinks on one occasion
- Positive urine test for drugs of abuse at any time during the study
- Current diagnosis of substance dependence (other than alcohol and nicotine)
- lifetime history of attempted suicide or self-injury; lifetime diagnosis of schizophrenia, bipolar disorder, obsessive compulsive disorder and/or eating disorder; current diagnosis of clinically significant major depressive disorder and/or anxiety disorder
- Positive viral hepatitis or human immunodeficiency virus (HIV) test; Creatinine ≥ 2 mg/dl; current or prior history of clinically significant medical diseases (neurological, cardiovascular, respiratory, gastrointestinal, hepatic, renal, endocrine, reproductive)
- Females only: breast-feeding and/or positive urine pregnancy test at any time during the study
- Current or prior history of alcohol-induced flushing
- Specific exclusion criteria related to IV ghrelin administration:
  - History of adverse reaction or hypersensitivity to IV ghrelin
  - Blood triglycerides level > 350 mg/dL, body weight ≥ 120 kg, body mass index (BMI) ≥ 30 kg/m^2^, and/or diabetes mellitus
  - history of chronic inflammatory bowel diseases (*e.g*., Crohn’s disease, ulcerative colitis, celiac disease)
  - Resting systolic blood pressure < 100 mmHg at screening, and/or history of clinically significant hypotension (*e.g*., syncope episodes or fainting)
- Specific exclusion criteria related to magnetic resonance imaging (MRI) scan:
  - Left-handedness
  - History of claustrophobia
  - Metal contraindicated for MRI (*e.g*., implant, pacemaker, prosthesis, shrapnel, irremovable piercing)
- Medication use:
  - Using any medication known to inhibit or induce alcohol metabolizing enzymes within 4 weeks prior to the study, including (but not limited to): chlorzoxazone, isoniazid, metronidazole, disulfiram
  - Using any medication known to interact with alcohol within 2 weeks prior to the study, including (but not limited to): isosorbide, nitroglycerine, benzodiazepines, warfarin, antidepressants (amitriptyline, clomipramine, nefazodone), antidiabetic agents (glyburide, metformin, tolbutamide), H_2_-blockers (cimetidine, ranitidine), muscle relaxants, antiepileptic drugs (phenytoin, phenobarbital codeine), narcotics (darvocet, percocet, hydrocodone)
  - Using antihistamines (*e.g*., in cough and cold preparations), pain relievers, anti-inflammatory drugs (*e.g*., aspirin, ibuprofen, acetaminophen, celecoxib, naproxen) within 72 hours prior to each experimental session

**S3.B. Standardized Meal Options**

Participants could choose from 3 options of standardized meals (see below) and were instructed to eat everything served. All meals were individually adjusted for each participant’s calorie needs calculated using the Mifflin-St. Jeor equation [1] with an activity factor of 1.5. All meal options had the same macronutrient distribution of ~15% protein, ~30% fat, ~55% carbohydrate. The standardized snack included 37 reduced fat Cheez-It® crackers (150 kcal - 12% protein, 30% fat, 59% carbohydrate).

For timing of the standardized meals, see **Figure 1** (main manuscript).

*Standardized breakfast:*

- Option 1: Vegetable omelet + Toast with butter and jelly + Orange juice
- Option 2: Turkey bacon, egg and cheese breakfast sandwich + Orange slices + Apple juice
- Option 3: Oatmeal with walnuts, brown sugar and dried cranberries + Milk

*Standardized lunch:*

- Option 1: Turkey sandwich + Potato chips + Grapes + Cranberry juice
- Option 2: Roast beef sandwich + Potato chips + Grapes + Lemonade
- Option 3: Vegetable wrap with hummus and provolone cheese + Orange slices + Potato Chips

*Standardized dinner:*

- Option 1: Spaghetti with turkey meatballs and parmesan cheese + Dinner roll with butter + Side salad with Italian dressing + Pineapple tidbits
- Option 2: Chicken breast + Mashed potatoes with gravy + Green beans + Bread and butter + Fruit punch + Tropical fruit cocktail
- Option 3: Penne pasta with spaghetti sauce and cheese blend + Side salad + Bread and butter + Lemonade

**S3.C. Blood Collection, Processing, and Assay**

Blood draws for both IV-alcohol experiments were done by inserting a 20 ga IV catheter into the peripheral vein of the left arm for multiple blood draws throughout the experiment. Plasma acyl-ghrelin was measured according to the same procedures listed in **S1.C**. As with other studies, plasma total-ghrelin was measured using commercially available ELISA kits (EMD Millipore Corp., Billerica, MA – Cat.# EZGRT-89K) following manufacturer instructions. The optical density of each plate was determined using SpectraMax M2 microplate reader (Molecular Devices, Sunnyvale, CA) in accordance with manufacturer’s instructions. A four or five parameter logistic regression was applied for quantification using Graph Pad Prism V.7 software (Graph Pad Software Inc., La Jolla, CA). Samples outside the quantification range of the calibration were diluted with assay buffer in accordance with protocol and re-measured. All measurements were carried out in duplicate.

**Appendix S4. *GHSR, GHRL,* and *MBOAT4* Expression in Human Post-Mortem Brain Tissue**

*RNA Extraction:*

Total RNA was extracted from male post-mortem frozen brain tissue, using RNeasy Lipid Tissue Mini kit (Qiagen, Hilden, Germany), and nucleic acid extraction was performed according to manufacturer’s protocols (<http://www.qiagen.com>). Tissue samples were manually disrupted using a rotary homogenizer and homogenate was passed through a RNeasy mini column containing RNeasy silica gel membrane, which selectively binds RNA. Total RNA was subsequently resuspended in RNAse-free water. RNA concentration and quality were determined using an Agilent 2100 Bioanalyzer and an RNA 6000 Nano kit. RNA samples were stored at -80°C.

*Real time qPCR and Data Analaysis:*

1 μg total RNA was reverse-transcribed using SuperScript ™ III First-Strand Synthesis Supermix for qRT-PCR kit (Invitrogen, Waltham, MA). Reverse transcription reactions were incubated at 25°C for 10 min, 50°C for 30 min, and 85°C for 5 min. Real-time quantitative PCR was run in ViiA™ 7 Real-Time PCR System using TaqMan Gene Expression Assays (Thermo Fisher, Waltham, MA- GHSR, Hs00269780_s1; GHRL, Hs01074053_m1; MBOAT4, Hs01074954_s1) The GADPH gene was used as an endogenous control. Data were analyzed using ViiA™ 7 Software (Applied Biosystems, Foster City, CA). Automatic cycle threshold (Ct) detection was normalized with the endogenous GAPDH gene by subtracting the Ct for *GAPDH* from the Ct for *GHSR, MBOAT4,* and *GHRL* mRNA for each brain region in each participant, yielding a ΔCt.

**Appendix S5. Effects of Ethanol on Ghrelin Levels in *Ghsr* KO and WT Rats**

For all animal experiments, whole blood was collected into microtubes treated with EDTA (1.7 mg/ml final concentration) and Pefabloc^®^ SC (Cat #11429876001, Millipore Sigma, Billerica, MA; 1 mg/mL final concentration). Samples were immediately centrifuged at 2,000 x g for 15 min at 4°C. Plasma was extracted and treated with 5% (v/v) 1 M HCl before storage at -80°C until assay. Acyl-ghrelin (AG) and des-acyl ghrelin (DAG) were measured using commercial ELISA kits purchased from ALPCO (AG: cat#-32-5117, DAG: cat#-32-5118, Salem, NH). Samples were measured in duplicate. The optical density of each plate was determined using a Bio-Rad Microplate Reader (Model#680; Bio-Rad Laboratories, Hercules, CA, USA). A 4 or 5 parameter logistic equation was applied for quantification using Bio-Rad Microplate Manager® Software.

**Appendix S6. Gastric Mucosal Cell Experiment**

Briefly, stomachs were isolated from chloral hydrate (700 mg/kg i.p.) anesthetized mice, inverted inside-out, cleared of their contents, and digested for 1.5 hrs at 37°C in 35 U dispase II/15 mL PBS (Roche Diagnostics, Indianapolis, IN). The mucosal cells were then scraped off the stomach into sterile DMEM/F-12 (1:1) medium (Mediatech Inc. Manassas, VA) containing 10% (vol/vol) FBS (Atlanta Biologicals, Lawrenceville, GA), supplemented with 100 U/mL penicillin and 100 μg/mL streptomycin sulfate. After centrifugation at 310 g for 3 min to remove the medium, the cells were digested with 0.25% trypsin EDTA (Mediatech Inc. Manassas, VA) for 5 min. The trypsin was inactivated with addition of FBS containing DMEM/F-12 medium, and the mucosal cells were dispersed mechanically using fire-polished pasteur pipettes. The cells were filtered through a 100 µM Falcon nylon mesh filter, and then centrifuged at 310 g for 3 min to remove the supernatant medium. The cells were resuspended in FBS containing DMEM/F-12 medium, plated at a density of 1 X 10^5^ cells/mL/well, and supplemented with sodium octanoate-bovine serum albumin (BSA) to achieve a final concentration of 50 μM sodium octanoate in poly-D-lysine pre-coated 24-well plates. The cells were incubated overnight in a 5% CO_2_ incubator at 37°C. The following day, 50X ethanol stock solutions were prepared by diluting 200 proof ethanol in ultrapure water. The cells were treated with medium containing different ethanol concentrations by addition of the stock solutions to serum free DMEM (Life Technologies, Grand Island, NY) containing 50 μM sodium octanoate-BSA and either 0 mM or 5 mM glucose. After incubating the cells for 6 h, the medium was collected, placed on ice and immediately centrifuged at 4°C at 800 g x 5 min. Hydrochloric acid was added to the supernatant to achieve a final concentration of 0.1 N (for stabilization of acyl-ghrelin) and stored at -80°C until analysis. Acyl-ghrelin concentration in the supernatants were measured by using ELISA kits (EZRGRA-90K; Millipore-Merck, St. Charles, MO). The endpoint calorimetric measurements for the ELISA were performed using a PowerWave XS2 Microplate spectrophotometer and Gen5 software (BioTek Instruments, Inc. Winooski, VT).

**Appendix S7. Effects of Ethanol on hGOAT activity**

Methyl arachidonyl fluorophosphonate (MAFP, Cayman Chemical, Ann Arbor, MI) was diluted in DMSO from a stock in methyl acetate. Absolute ethanol was purchased from Pharmco (Brookfield, CT). Octanoyl-coenzyme A (Octanoyl-CoA, CoALA Biosciences, Austin, TX) was diluted to 5 mM in 10 mM Tris-HCl pH 7.0 and stored in low adhesion tubes at -80°C until use. The ghrelin-mimetic GSSFLC_NH2_ peptide substrate was commercially synthesized by Sigma-Genosys (The Woodlands, TX) in Pepscreen format. The GSSFLC_NH2_ peptide was solubilized in 1:1 acetonitrile/water solution and stored at -80°C. The peptide concentration was determined spectrophotometrically at 412 nm by reaction of the cysteine thiol with 5,5’dithiobis (2-nitrobenzoic acid) using an ϵ_412_ of 14,150 M^-1^ cm^-1^ [2]. Peptide substrates were fluorescently labeled with acrylodan and purified via reverse phase HPLC, as previously reported. hGOAT was expressed and enriched in insect (Sf9) membrane protein fractions using a previously published procedure [3, 4].

Membrane protein fractions from Sf9 cells expressing hGOAT were thawed on ice and then homogenized by passage through an 18-gauge needle ten times. Ethanol working stocks (5.44 mM, 10.87 mM, 21.74 mM, 43.48 mM, 65.22 mM, and 86.96 mM) were prepared by serial dilution. Assays were performed with 70 µg of membrane protein as determined by Bradford assay. For each set of eleven assays, a master mix (495 µL) was prepared containing 2.5 mM HEPES, 10 µM methyl arachidonyl fluorophosphonate (MAFP), and 770 µg membrane protein. The master mix was aliquoted (44 µL) into each low-adhesion microcentrifuge tube, followed by addition of 1 µL of the appropriate ethanol stock. The reaction mixture was then pre-incubated for 30 min at room temperature in a sealed microcentrifuge tube. Reactions were initiated by the addition of octanoyl-CoA (300 µM final concentration) and GSSFLC_AcDan_ peptide (1.5 µM final concentration) to yield a final reaction volume of 50 µL. Reactions were incubated for 1 h at room temperature while sealed, and then stopped by addition of 50 µL of 20% acetic acid in isopropanol. Membrane proteins were then removed via precipitation with 16.7 µL of 20% trichloroacetic acid followed by centrifugation (1000 x *g*, 1 min). The supernatant was then analyzed via reverse-phase HPLC, as described previously [5, 6]. GOAT acylation activity was determined by substrate and product peak integration in the presence of either ethanol or water (vehicle). Percent activity for each reaction was calculated using equations 1 and 2 [3].

$$\left( \text{1} \right)\text{ \% peptide octanoylation = }\frac{\text{fluorescence of octanoylated peptide}}{\text{total peptide fluorescence (substrate and product)}}$$

$$\left( \text{2} \right)\text{ \% activity}\text{ }\text{=} \frac{\text{\% peptide octanoylation in presence of inhibitor}}{\text{\% peptide octanoylation in absence of inhibitor}}$$

**Table S1: Baseline Characteristics of the Samples Enrolled in Human Laboratory Experiments**

|  | **Oral Priming & ASA** | **Oral Fixed Dose** | **IV ASA** | **IV Fixed Dose** |
| --- | --- | --- | --- | --- |
| Sample Size | 16 | 12 | 11 | 6 |
| Gender (n, Male%) | 13, 81.3% | 11, 91.7% | 8, 72.7% | 5, 83.3% |
| Race (n, Black/African American%) | 12, 75% | 11, 91.7% | 9, 81.8% | 5, 83.3% |
| Age (years, M±SD) | 42 ± 10.8 | 40.5 ± 13.1 | 39.7 ± 11.7 | 39.0 ± 7.6 |
| BMI (kg/m^2^, M±SD) | 27.9 ± 4.1 | 26.9 ± 4.7 | 25.9 ± 2.8 | 25.3 ± 2.8 |
| Current Tobacco Smoker  (n, Smoker%) | 7, 43.8% | 9, 75% | 8, 72.7% | 5, 83.3% |
| 90-day DDD (M±SD)^a^ | 9.1 ± 6.5 | 11.3 ± 4.7 | 8.8 ± 7.3 | 10.1 ± 6.8 |

^a^ Determined by Timeline Followback (TLFB) at screening visit

IV = Intravenous, ASA = Alcohol Self-Administration, BMI = Body Mass Index, DDD = drinks per drinking day

**Table S2: Additional Information about the Human Laboratory Experiments**

|  | **IV Fixed Dose** | **IV ASA** | **Oral Fixed Dose** | **Oral Priming & ASA** |
| --- | --- | --- | --- | --- |
| Meal Status at Beginning of Alochol Administration Session (minutes post-meal) | +80 | +45 | +120 | +175 |
| Alcohol Type | 6.0% (v/v) Ethanol | 6.0% (v/v) Ethanol | 40.0% (v/v) Liquor  (80 proof Smirnoff Vodka) | Preferred Alcohol ^a^ |
| Alcohol Diluent | Saline | Saline | Preferred Mixer^b^ | Preferred Mixer^b^  (only if liquor selected) |
| Target BAC (mg%) | 0.08 | Limit – 0.12  (0.0075% per infusion) | 0.06 | Limit – 0.12  (P: 0.03/drink, ASA:0.015 /drink) |
| Total Duration of Alcohol Administration (min) | 35 | 120 | 20 | 135  (P – 15, ASA – 120) |
| Total Alcohol Administered  (g, M ± SD) | 31.3 ± 9.4 | 27.2 ± 15.2 | 33.4 ± 8.1 | 56.4 ± 25.3 |
| Time of Baseline Blood Draw  (min post-meal, min pre-alcohol) | 12:45 PM  (+35, -45) | 11:35 AM  (+15, -30) | 8:30 AM  (+60, -90) | 8:30 AM  (-30, -280) |
| Baseline Acyl-Ghrelin Concentration  (pg/mL, M±SD) | 89.4 ± 15.9 | 71.1 ± 32.9 | 52.9 ± 22.6 | 71 ± 31.4 |
| Baseline Total Ghrelin Concentration  (pg/mL, M ± SD) | 475.7 ± 106.1 | 519.1 ± 208.4 | 381.5 ± 104.0 | 417.2 ± 235.9 |

^a^ No limitations placed on type or brand of alcohol able to be selected. ^b^See S1C for preferred mixer options able to be chosen.

IV = intravenous, ASA = alcohol self-administration, BAC = blood alcohol concentration, P = priming

**Table S3: Baseline Demographics of the Sample Enrolled in the Human Post-Mortem Study**

|  | **AUD** | **Control** |
| --- | --- | --- |
| Sample Size (N) | 11 | 16 |
| Age (years, Mean ± SD) | 50.5 ± 6.1 | 49.9 ± 11.3 |
| BMI (kg/m^2^, Mean ± SD) | 24.6 ± 5.4 | 33.1 ± 10.5 |
| Smoker (N, %) | 11, 100% | 2, 15.4% |
| Pack Years Smoking (years, Mean ± SD) | 45.0 ± 19.2 | 4.1 ± 13.9 |
| Brain pH (Mean ± SD) | 6.6 ± 0.4 | 6.7± 0.2 |
| PMI (hrs, Mean ± SD) | 38.9 ± 12.5 | 31.1 ± 13.9 |
| ^a^Cause of Death (N, % Cardiac) | 4, 36.4% | 13, 81.3% |

^a^ Causes of death among controls: cardiac, respiratory, and unknown. Causes of death among AUD: cardiac, respiratory, hepatic, toxicity, blood loss, vascular, and unknown

AUD = Alcohol Use Disorder, BMI = Body Mass Index, PMI = postmortem interval

**Figure S1: Flow Diagram of Inclusion in Oral Alcohol Priming and Alcohol Self-Administration Analysis**

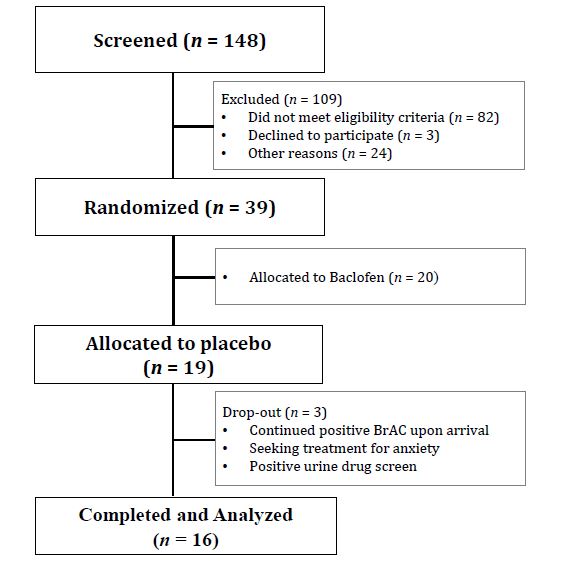

**Figure S2: Flow Diagram of Inclusion in Oral Fixed Alcohol Administration Analysis**

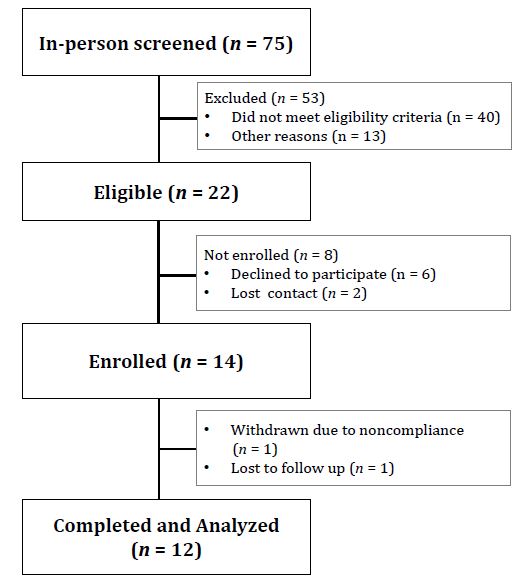

**S3: Flow Diagram of Inclusion in Intravenous Alcohol Self-Administration Experiment**

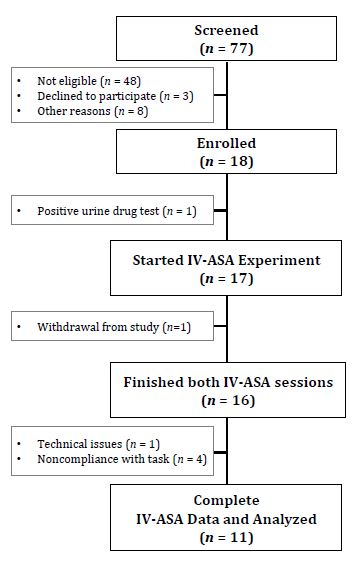

**Figure S4**: Flow Diagram of Inclusion in Fixed Intravenous Alcohol Administration Experiment

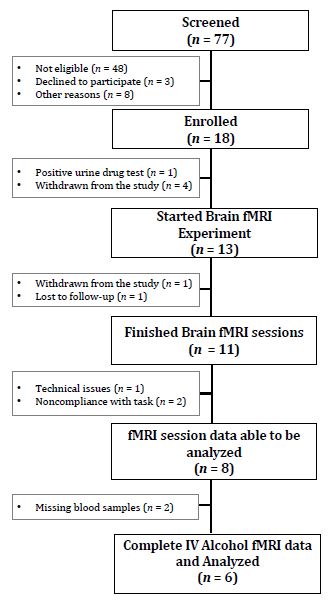

**Figure S5:** **Effect of Ethanol on hGOAT Activity**
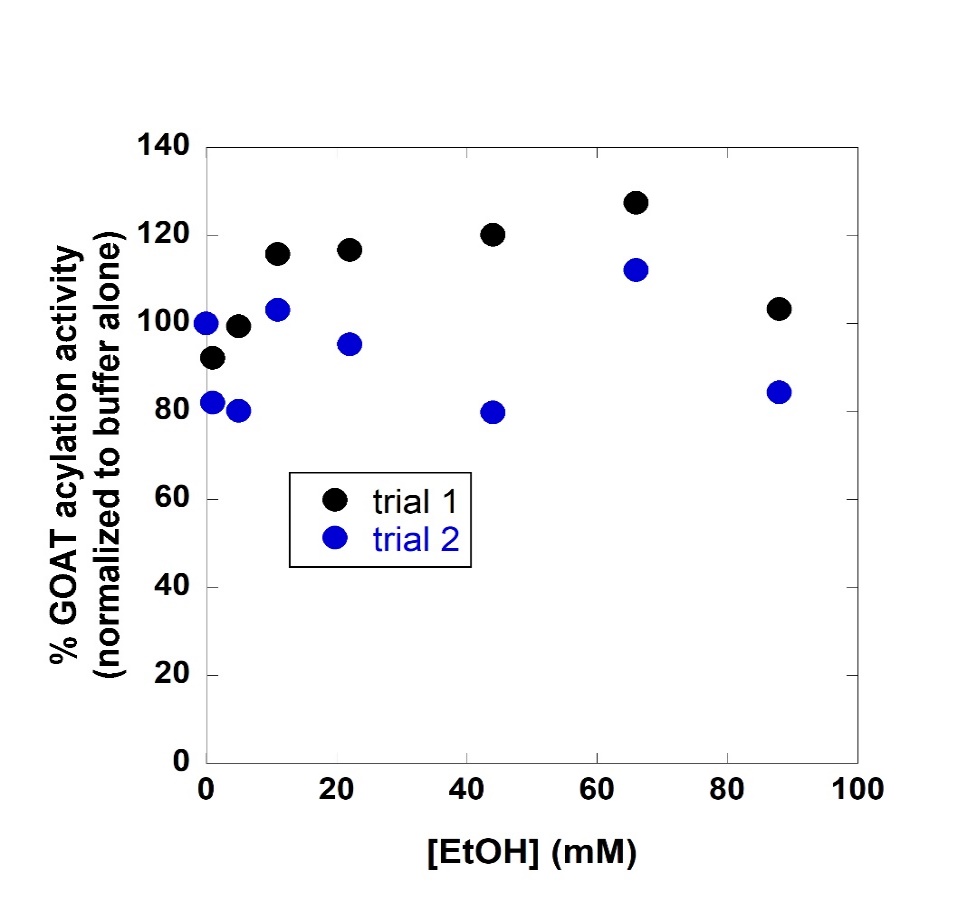

GOAT = Ghrelin O-acyl transferase; EtOH = ethanol

Trial 1 and Trial 2 are data from replications of the same expeirment

**Figure S6: Effect of Ethanol and Sucrose on AG:DAG Ratio in Rats**

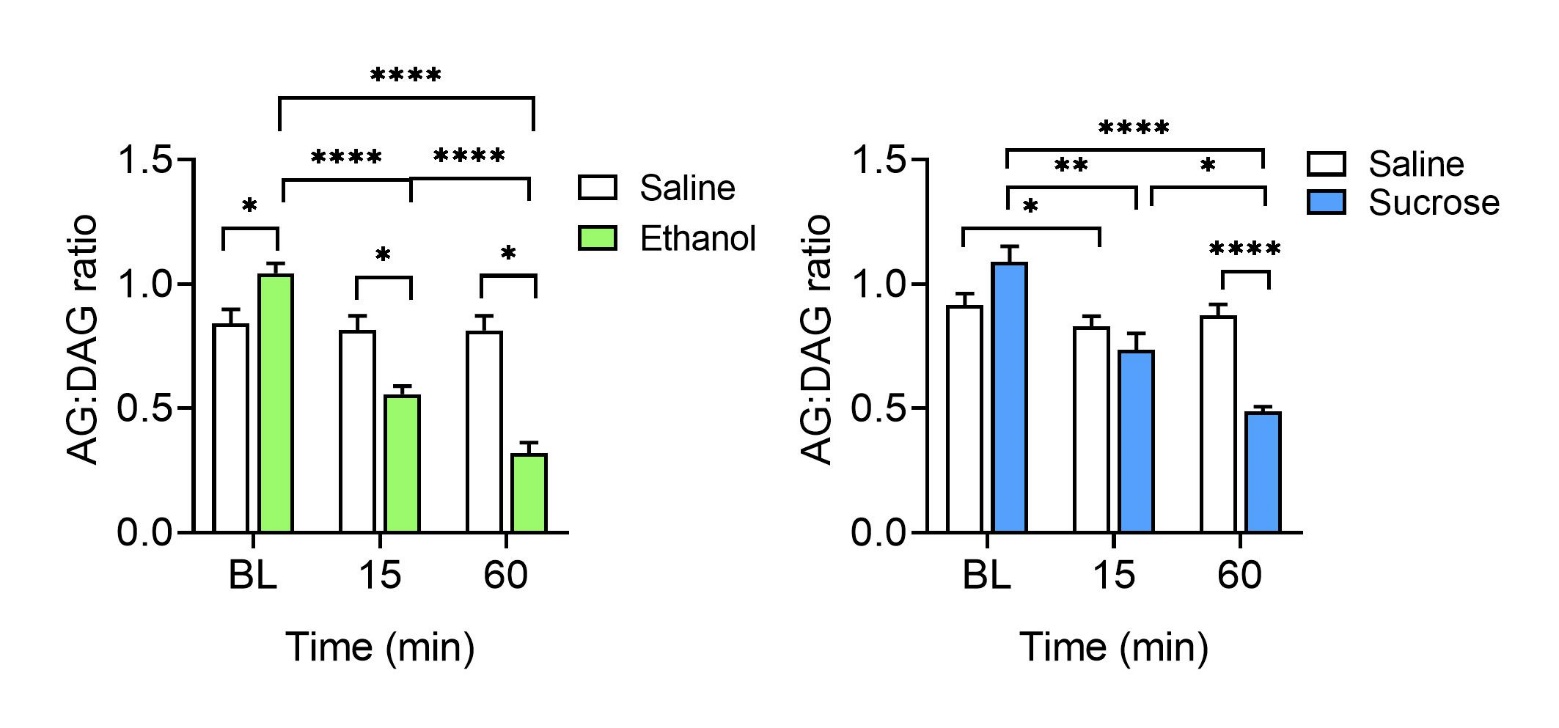

AG:DAG = acyl-ghrelin:des-acyl-ghrelin, BL = baseline (0 min)

Data represents AG:DAG ratio after ethanol treatment (left, green) vs. saline (white) and after sucrose treatment (right, blue) vs. saline (white) in male Wistar rats. *Ethanol (left):* Significant main effect of treatment [F(1, 18) = 10.68, p = 0.0043], time [F(2, 36) = 59.03, p < 0.0001], and interaction [F(2, 36) = 50.02, p < 0.0001]. Post-hoc testing revealed that AG:DAG ratio was significantly lower following ethanol treatment at 15 min (p < 0.0001) and 60 min (p < 0.0001), compared to baseline, as well as at 60 min, compared to 15 min (p < 0.0001). In comparison to saline treatment, ethanol treatment had a significantly lower AG:DAG ratio at BL (p = 0.0168), 15 min (p = 0.0014), and 60 min (p < 0.0001). *Sucrose (right):* Significant main effect of treatment [F(1, 16) = 4.93, p = 0.041], time [F (1.966, 31.45) = 26.38, p < 0.0001], and treatment × time interaction [F (2, 32) = 19.18, p < 0.0001]. Post-hoc testing revealed that sucrose significantly lowered the AG:DAG ratio at both 15 min (p = 0.007) and 60 min (p < 0.0001) relative to baseline and at 60 min relative to 15 min (p = 0.0331). Rhe AG:DAG ratio was significantly lower at 60 min following sucrose treatment vs. saline treatment (p = 0.013). Post-hoc Sidak’s multiple comparison tests: *p < 0.05, **p < 0.01, ***p < 0.001, ****p < 0.0001
