## Supplementary material for "Pursuing the Mechanisms Underlying Alcohol-Induced Changes in the Ghrelin System: New Insights from Preclinical and Clinical Investigations": NIH Cover Sheet

### NIH Publishing Agreement & Manuscript *Cover Sheet*

By signing this Cover Sheet, the Author, on behalf of NIH, agrees to the provisions set out below, which **modify and supersede,** solely with respect to NIH, any conflicting provisions that are in the Publisher's standard copyright agreement (the "Publisher's Agreement"). If a Publisher's Agreement is attached, execution of this Cover Sheet constitutes an execution of the Publisher's Agreement, subject to the provisions and conditions of this Cover Sheet.

1. **Indemnification.** No Indemnification or "hold harmless" obligation is provided by either party.

2. **Governing Law.** This agreement will be governed by the law of the court in which a claim is brought.

3. **Copyright.** Author's contribution to the Work was done as part of the Author's official duties as a NIH employee and is a Work of the United States Government. Therefore, copyright may not be established in the United States. 17 U.S.C. § 105. If Publisher intends to disseminate the Work outside of the U.S., Publisher may secure copyright to the extent authorized under the domestic laws of the relevant country, subject to a paid-up, nonexclusive, irrevocable worldwide license to the United States in such copyrighted work to reproduce, prepare derivative works, distribute copies to the public and perform publicly and display publicly the work, and to permit others to do so.

4. **No Compensation.** No royalty income or other compensation may be accepted for work done as part of official duties. The author may accept for the agency a limited number of reprints or copies of the publication.

5. **NIH Representations.** NIH represents to the Publisher that the Author is the sole author of the Author's contribution to the Work and that NIH is the owner of the rights that are the subject of this agreement; that the Work is an original work and has not previously been published in any form anywhere in the world; that to the best of NIH's knowledge the Work is not a violation of any existing copyright, moral right, database right, or of any right of privacy or other intellectual property, personal, proprietary or statutory right; that where the Author is responsible for obtaining permissions or assisting the Publishers in obtaining permissions for the use of third party material, all relevant permissions and information have been secured; and that the Work contains nothing misleading, obscene, libelous or defamatory or otherwise unlawful. NIH agrees to reasonable instructions or requirements regarding submission procedures or author communications, and reasonable ethics or conflict of interest disclosure requirements unless they conflict with the provisions of this Cover Sheet.

6. **Disclaimer.** NIH and the Author expressly disclaim any obligation in Publisher's Agreement that is not consistent with the Author's official duties or the NIH mission, described at http://www.nih.gov/about/. NIH and the Author do not disclaim obligations to comply with a Publisher's conflict of interest policy so long as, and to the extent that, such policy is consistent with NIH's own conflict of interest policies.

7. **For Peer-Reviewed Papers to be Submitted to PubMed Central.** The Author is a US government employee who must comply with the NIH Public Access Policy, and the Author or NIH will deposit, or have deposited, in NIH's PubMed Central archive, an electronic version of the final, peer-reviewed manuscript upon acceptance for publication, to be made publicly available no later than 12 months after the official date of publication. The Author and NIH agree (notwithstanding Paragraph 3 above) to follow the manuscript deposition procedures (including the relevant embargo period, if any) of the publisher so long as they are consistent with the NIH Public Access Policy.

8. **Modifications.** PubMed Central may tag or modify the work consistent with its customary practices and with the meaning and integrity of the underlying work.

The NIH Deputy Director for Intramural Research, Michael Gottesman, M.D., approves this publishing agreement and maintains a single, signed copy of this text for all works published by NIH employees, and contractors and trainees who are working at the NIH. No additional signature from Dr. Gottesman is needed.

Author's name: Lorenzo Leggio

Author's Institute or Center: NIDA Check if Publisher' Agreement is attached ☐

Name of manuscript/work: Pursuing the Mechanisms Underlying Alcohol-Induced Changes in the Ghrelin Sy

Name of publication: bioArxiv

Lorenzo Leggio -SR1 Digitally signed by Lorenzo Leggio -SR1 Date: 2020.03.22 10:50:27 -04'00'

Author's signature

07/29/2020

Date

NIH OGC August 2008
